# Monolith, porous bead, membrane or fibre? A comparative characterisation of four oligo-dT affinity chromatography matrices for mRNA capture

**DOI:** 10.64898/2026.11.17.767441

**Authors:** Jixin Qu, Manoj Pohare, Shady Hamed, Adithya Nair, Mabrouka Maamra, Zoltán Kis

**Affiliations:** School of Chemical, Materials and Biological Engineering, University of Sheffield, Sir Robert Hadfield Building, Mappin Street, Sheffield S1 3JD, UK; Department of Chemical Engineering, Imperial College London, Roderic Hill Building, South Kensington Campus, London SW7 2AZ, UK; Downstream Processing Group, Bioprocessing Technology Institute (BTI), Agency for Science, Technology and Research (A*STAR), 20 Biopolis Way, #06-01 Centros, 138668, Singapore

**Author notes:** corresponding author: Manoj Pohare,; Zoltan Kis. These authors contributed equally (Jixin Qu, Manoj Pohare).

**Keywords:** oligo-dT affinity chromatography, monolith chromatography, bead chromatography, membrane chromatography, fibre adsorber, design of experiments, process economics, mRNA purification

## Abstract

Oligo-deoxythymidine (oligo-dT) affinity chromatography is the most widely used capture step for *in vitro* transcribed mRNA and is now offered in four stationary-phase architectures, yet, to our knowledge, no published study has compared all four. We characterised a monolith, perfusive porous beads, a membrane adsorber and an electrospun cellulose fibre adsorber with a short (eGFP, 930 nt) and a long (SARS-CoV-2 spike, S-2P, 4284 nt) mRNA. After screening the loading salt (NaCl or guanidine hydrochloride, Gu-HCl), we mapped load flow rate, load concentration and Gu-HCl concentration with response-surface designs and compared the formats at near-optimal conditions. Gu-HCl gave a higher peak recovery than NaCl in seven of the eight matrix–construct combinations. At the near-optimal runs, the fibre adsorber recovered 17.9 mg eGFP and 13.3 mg S-2P per mL of bed, 2.3-to 2.6-fold the membrane and 3.5-to 4.8-fold the monolith and beads (p < 0.001); these are yields at loads that differed between formats (5 to 20 mg/mL of bed), not binding capacities. Productivity ranked fibre > membrane > monolith > beads (0.314 to 0.035 mg/mL/min for eGFP). The membrane and fibre adsorber lost yield sharply above 0.4–0.6 M Gu-HCl, whereas the monolith and beads declined gradually. In an amortised cost model at baseline cost inputs, the fibre adsorber was the least expensive format per milligram of purified mRNA at every number of device cycles (USD 114–140/mg for single use and 78–92/mg at 20 cycles), and the membrane the most expensive. A one-at-a-time sensitivity analysis over wide ranges of the mRNA feed cost, the labour and facility rate, the device price and the number of cycles per device left the fibre adsorber the least expensive format in almost every combination examined, with the mRNA feed cost the dominant input. Only the monolith and beads are currently marketed as GMP-grade, process-scale materials, so the choice of format depends on whether capacity, productivity, cost or manufacturing readiness takes priority.

## 1. Introduction

Messenger RNA (mRNA) is now an established vaccine modality and is expanding rapidly into protein replacement, gene editing and immuno-oncology [1,2]. Its manufacturing platform is largely product-agnostic: a linearised plasmid template is transcribed *in vitro* by T7 RNA polymerase. The resulting crude *in vitro* transcription (IVT) reaction, which contains the mRNA together with residual nucleoside triphosphates (NTPs), enzymes, template DNA, abortive transcripts and double-stranded RNA (dsRNA), is purified before formulation [3–5]. Whereas IVT is fast and productive, downstream purification has become the throughput-and cost-determining part of the drug-substance process, and the removal of dsRNA and other impurities during purification is directly linked to the safety and potency of the product [5,6].

Oligo-deoxythymidine (oligo-dT) affinity chromatography is the most widely used capture step for IVT mRNA because it is selective for polyadenylated species, scalable and compatible with the salt-containing IVT matrix [5,7]. It relies on Watson–Crick hybridisation between the 3′ poly(A) tail of the mRNA and oligo-dT ligands immobilised on the stationary phase. Each adenine–thymine pair contributes two hydrogen bonds; individually weak, they collectively confer high stability and specificity on the poly(A)–oligo-dT duplex, whose free energy is further governed by base stacking, counter-ion screening of the two polyanionic backbones and the loss of conformational entropy that opposes binding: in solution the poly(A) is flexible and samples many conformations, whereas once hybridised to a surface-bound ligand it is held in a single, comparatively rigid duplex [8–10]. Duplex stability is usually predicted with the nearest-neighbour model, which sums experimentally determined free-energy contributions for each pair of adjacent base pairs. Because the poly(A)-oligo-dT duplex is an RNA/DNA hybrid, whose stability differs from that of the corresponding DNA duplex, nearest-neighbour parameters for RNA/DNA hybrids, rather than those for DNA duplexes, are the appropriate thermodynamic reference [11,12]. The capacity and yield of an oligo-dT step therefore depend on the ionic strength and nature of the loading salt (sodium chloride, or the chaotrope guanidine hydrochloride, Gu-HCl, which also relaxes mRNA secondary structure and exposes the poly(A) tail) [7,13,14], on the residence time available for the tail to find and align with a ligand [15,16], and on the architecture and ligand chemistry of the matrix [15,17,18].

Four architectures of oligo-dT stationary phase are now commercially available, and each resolves the trade-off between accessible surface area, mass-transfer resistance, residence time, pressure limits and ligand density differently. Porous beads packed into columns are the historical format: the ligand is presented inside a pore network that mRNA must enter by diffusion, a limitation that perfusive beads with large through-pores reduce but do not remove [19]. Monoliths present the ligand on the walls of continuous macroporous channels, so that transport is predominantly convective and capacity is largely independent of flow rate [8]. Membrane adsorbers are thin, high-surface-area polymer sheets operated at residence times of seconds [20]. Most recently, electrospun cellulose nanofibre adsorbers, developed for high-productivity, rapid-cycling protein capture [21], have been functionalised with oligo-dT and commercialised as Fibro dT [22]. In prototype form, this adsorber showed dynamic binding capacities about 2–4-fold higher than a monolith and 2–13-fold higher than porous beads [15]. Which format best suits a given mRNA product and manufacturing scenario is not obvious a priori. Direct comparisons remain scarce, and to our knowledge none has covered all four formats or extended beyond binding capacity to productivity and process economics.

Here we characterised one representative small-scale column of each format, a 1 mL CIMmultus Oligo-dT 18 monolith, a 1 mL POROS GoPure Oligo (dT)25 beads column, a 0.2 mL Purexa OdT Maxi membrane and a 0.4 mL Fibro dT fibre adsorber, with two model constructs spanning the size and poly(A) tail range of vaccine and therapeutic mRNAs: enhanced green fluorescent protein mRNA (eGFP, 930 nt, including a 49-nt poly(A) tail) and mRNA encoding SARS-CoV-2 spike glycoprotein, prefusion-stabilised by two proline substitutions (K986P/V987P; S-2P mRNA, 4284 nt, including a 120-nt split poly(A) tail). We screened loading salt type and concentration, mapped the yield as a function of load flow rate, load concentration and Gu-HCl concentration with three-factor response-surface designs and compared the formats at near optimal conditions in terms of yield, productivity, relative cost and manufacturing readiness. The resulting operating windows, interpreted through duplex thermodynamics and mass transfer, are intended to guide the selection and operation of oligo-dT capture steps in mRNA vaccine and therapeutic manufacturing.

## 2. Materials and methods

### 2.1 mRNA transcripts prepared by *in-vitro* transcription

The mRNA transcripts for eGFP (930 nt, including a 49-nt poly(A) tail), and S-2P (4284 nt, including a 120-nt split poly(A) tail), were synthesised using *in-vitro* transcription (IVT). The IVT reactions used linearised plasmid DNA templates for both eGFP and S-2P (GenScript, New Jersey, USA; the template DNA sequences are given in [23], where S-2P is referred to as CSP), and the following materials. T7 bacteriophage RNA polymerase, nucleoside triphosphates (ATP, CTP, GTP and UTP), HEPES buffer (pH 7.3), dithiothreitol (DTT), spermidine, inorganic pyrophosphatase and RNase inhibitor were obtained from Roche (Mannheim, Germany). Magnesium acetate and sodium chloride were supplied by Sigma-Aldrich (Gillingham, UK).

For IVT, the reaction components were assembled in a HEPES buffer, briefly vortexed and incubated at 37 °C for 120 min in a shaking incubator at 300 rpm as described in [24]. Following incubation, a 20 µL aliquot was diluted to 50 µL with nuclease-free water. Samples were purified by solid-phase extraction on silica spin columns (New England Biolabs) according to the manufacturer’s instructions, and the mRNA was eluted in 100 µL of nuclease-free water. mRNA concentration was determined from the absorbance at 260 nm using a NanoDrop™ One^C^ UV-Vis spectrophotometer (Thermo Fisher Scientific), normalised to a 1.0 cm path length.

### 2.2 Chromatography equipment, columns, buffers and load

All chromatography experiments were conducted using the ÄKTA Pure 25 chromatography system (Cytiva, Uppsala, Sweden). This system was configured with a Micro Kit, two independent binary pumps (A and B) connected to four inlet lines (A1, A2, B1 and B2), a multi-wavelength UV detector operating at 260 nm, and an inline conductivity monitor. Samples were loaded onto the column using a 10 or 50 mL Superloop, and for sample volumes exceeding this capacity, multiple loading cycles were performed. All runs were performed at room temperature between 20-25°C. Chromatographic methods were developed and executed, and data were processed, using UNICORN software (Version 7.8).

The four oligo-dT columns utilised in this study are summarised in Table 1, including a 1 mL CIMmultus® Oligo-dT 18 monolith column (Sartorius BIA Separations, Ajdovščina, Slovenia), a 1 mL POROS^TM^ GoPure Oligo (dT)25 beads column (Thermo Fisher Scientific, MA, USA), a 0.2 mL Purexa^TM^ OdT Maxi membrane column (Donaldson Purilogics, SC, USA), and a 0.4 mL Fibro dT column (Cytiva, Uppsala, Sweden). The standard oligo-dT chromatography process comprised five phases (Table S1): equilibration, sample loading, washing, elution, and cleaning-in-place (CIP). During the equilibration step, the column was conditioned with an equilibration buffer to establish optimal binding conditions for mRNA. The sample containing mRNA was then loaded onto the column, allowing selective binding. This was followed by a wash step to remove non-specifically bound impurities. Finally, the target mRNA was eluted with isocratic elution by flushing the column with nuclease-free water, and then the column was subsequently cleaned for reuse in the CIP phase. A single unit of each column was used for all experiments with that format, and no significant drift was noticed among the runs.

**Table 1.** Properties of four oligo-dT columns, as specified by the suppliers [. **15,22,25–27]**

| **Column type** | **Monolith** | **Beads** | **Membrane** | **Fibre** |
|---|---|---|---|---|
| Product | CIMmultus Oligo-dT 18 | POROS GoPure Oligo (dT)25 | Purexa OdT Maxi | Fibro dT adsorber |
| Base matrix | Poly(methacrylate) monolith, convective 2 µm macropores | 50 µm perfusive polystyrene-divinylbenzene beads | Polymer membrane, 0.45 µm pore size | Electrospun cellulose fibres with 0.7 µm pore size |
| Oligo-dT ligand | 18-mer, C12 spacer | 25-mer, C12 spacer | 25-mer, C12 spacer | Proprietary length (prototype 20-mer), single-point 5′ attachment, >6 µmol/g |
| Bed volume [mL] | 1 | 1 | 0.2 | 0.4 |

All buffers for the oligo-dT chromatography experiments were freshly prepared using nuclease-free water that meets European Pharmacopoeia standards and reagents of molecular biology grade. The buffers included the equilibration and loading buffer, which consisted of 50 mM sodium phosphate (pH 7.2) (BioWorld, TX, USA), with varying concentrations of NaCl (0.2–1.2 M) (Sigma-Aldrich, Gillingham, UK) or Gu-HCl (0.1–1.0 M) (Thermo Fisher Scientific, Oxford, UK) and 5 mM EDTA (pH 8.0, Thermo Fisher Scientific, Oxford, UK), with the final buffer pH of 7.4 ± 0.2. The elution buffer used nuclease-free water, while a 0.1 M or 0.5 M NaOH solution (Sigma-Aldrich, Gillingham, UK) was employed as the cleaning-in-place (CIP) solution according to the column manufacturer’s instructions. All buffers underwent degassing using an Elmasonic Easy ultrasonic cleaning device (Elma, Singen, Germany) by sonication at room temperature for 30 min to remove dissolved gases.

For column loading, the mRNA concentration of the crude IVT reaction mixture (not treated with DNase) was first determined by A260 after clean-up of an aliquot on a silica spin column (Section 2.1). The crude IVT mixture was then diluted to the target mRNA concentration in an equilibration buffer (50 mM sodium phosphate, 5 mM EDTA) containing Gu-HCl or NaCl at the concentration set by the experimental design. This load was then incubated at 50 °C for 5 min, and applied to the column. For the monolith, beads and membrane, the mass of mRNA loaded was fixed in advance, decided based on preliminary experimental data, which was just enough to see the breakthrough coming out from the column, whereas the fibre adsorber was loaded until breakthrough was observed in the flow-through UV signal or until the column became blocked, which occurred particularly at higher salt and chaotrope concentrations (Figure S1).

### 2.3 mRNA quantification

The mRNA yield was quantified by using a NanoDrop^TM^ One^C^ UV-vis spectrophotometer (Thermo Fisher Scientific, MA, USA). A 2 µL aliquot of each well-mixed sample was loaded onto the lower measurement pedestal, and the UV absorbance was recorded at 260 nm, with the instrument automatically adjusting the path length and normalising the reading to a 1.0 cm equivalent path length. Concentrations were calculated from the Beer-Lambert law using the standard mass extinction coefficient for single-stranded RNA (1.0 A260 unit = 40 µg/mL).

### 2.4 Experimental design and statistical analysis

#### 2.4.1 Salt screening

For each matrix-construct pair, the loading salt (NaCl, 0.2-1.2 M; Gu-HCl, 0.1–1.0 M) and its concentration were varied one factor at a time and the mass of mRNA recovered in the eluate was determined in triplicate (Figure 1, Table S2). The load mass in the salt screen was 5 mg for the monolith and the beads and 2.5 mg for the membrane (Table S2), so the membrane salt screen was run at a load challenge of 12.5 mg/mL of bed against 10 mg/mL in the response-surface designs; the fibre load varied with the salt concentration. In a few cases, higher Gu-HCl and NaCl concentrations were not tested. This was because, in the other transcript–matrix combinations, recovery never improved again once it had started to fall with increasing salt and chaotrope concentration.

**Figure 1.**
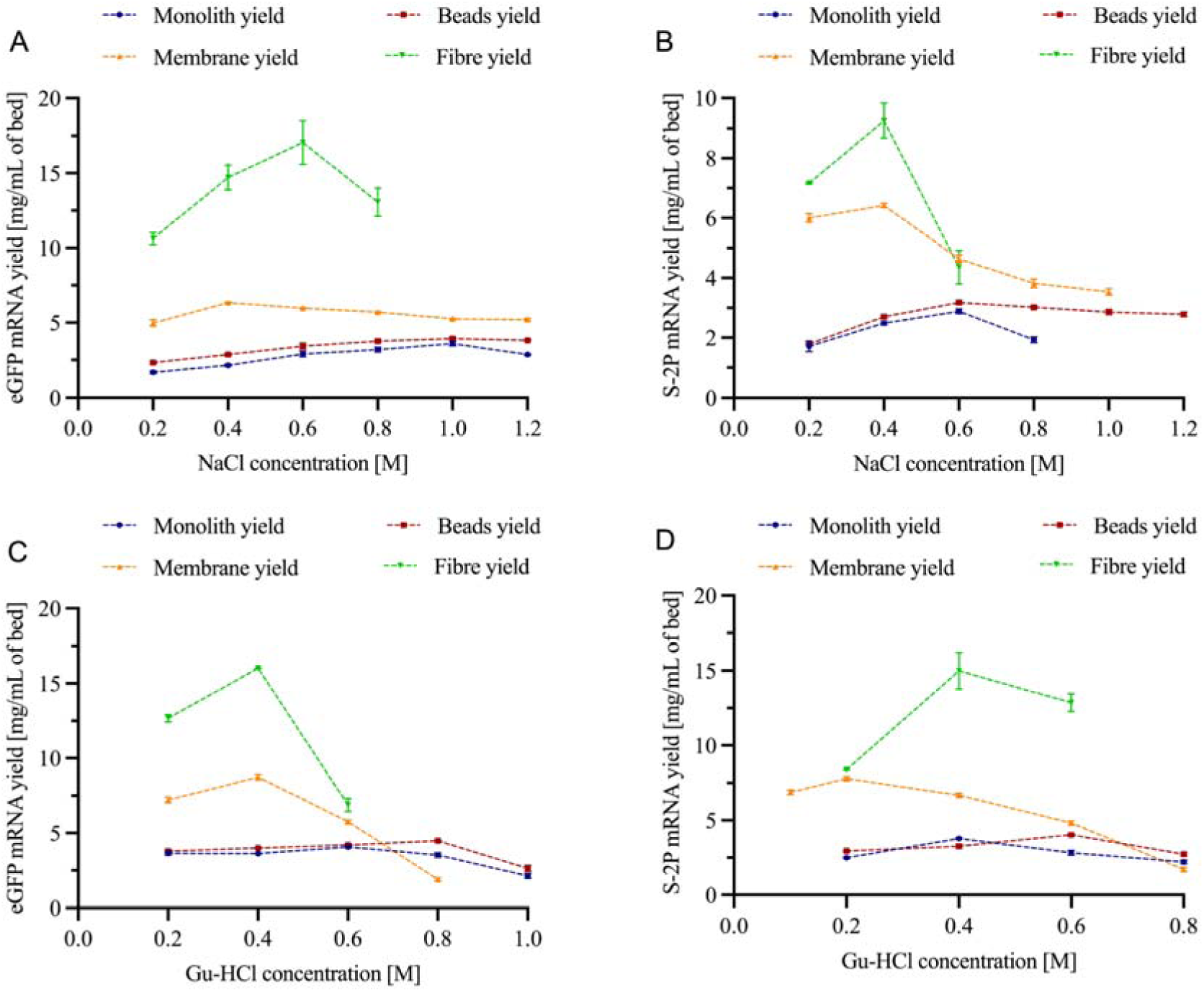
Yield of eGFP (A, C) and S-2P (B, D) mRNA per mL of bed on the four oligo-dT matrices as a function of NaCl (A, B) and Gu-HCl (C, D) concentration in the loading buffer. Symbols are means and error bars the standard deviation of three replicates. The fibre S-2P point at 0.6 M Gu-HCl (D) was not reproduced in the response-surface runs and was excluded from the response-surface model (Section 2.4.2).

#### 2.4.2 Response-surface designs

For each matrix-construct pair, the effects of Gu-HCl concentration, load flow rate and mRNA load concentration on the yield of mRNA per mL of bed were mapped with a three-factor, face-centred central composite response-surface design using JMP 19 (JMP Statistical Discovery LLC, Cary, NC, USA). The factor ranges for each format are given in Table 2. Each design comprised 16 runs (8 factorial, 6 axial and 2 centre points); the fibre designs were augmented to 20 runs with additional runs at 0.4 M Gu-HCl. Most runs were performed in duplicate or triplicate, and every replicate was entered into the analysis. Quadratic models (linear, interaction and squared terms) in coded factor levels (−1, 0, +1) were fitted by least squares to the replicate-level data and assessed by analysis of variance (ANOVA), the coefficient of determination (R^2^), adjusted R^2^, the predicted R² (from the PRESS statistic), the root-mean-square error (RMSE) and a lack-of-fit test against the pure error of replicate runs, and the model-predicted maximum within the design space was taken as the response-surface optimum [28]. The relative importance of each process parameter was expressed by its coded (standardised) coefficient and by the share of the model sum of squares attributable to its terms (Type III sums of squares; Tables S3, S4a and S4b). One fibre S-2P replicate (run 14, 3.40 mg against 1.04 and 0.82 mg) was excluded as an outlier, on the criterion that a replicate was set aside when it differed from the median of its own run by more than 100%; this criterion was applied to every run of every matrix-construct pair and excluded no other replicate. To avoid duplicating experiments, salt-screening runs performed at conditions close to fibre design points (0.2, 0.4 and 0.6 M Gu-HCl at 0.2 mg/mL) were incorporated into the fibre datasets (four runs per construct). These runs were performed at a load flow rate of 1.6 mL/min (Table S2) and were entered at the nearest design level, 1.5 mL/min; the difference corresponds to 0.05 in coded units and to less than 1 s of residence time, and flow rate was not a significant factor in either fibre model (Table S4a). Their consistency with the response-surface runs was assessed against the 95% prediction interval, for the mean of the same number of replicates, of a model fitted to the response-surface runs alone (Table S4c). Seven of the eight incorporated runs fell within this interval. The exception was fibre S-2P run 17 (0.6 M Gu-HCl, 0.2 mg/mL, 1.5 mL/min; 12.85 ± 0.59 mg/mL, n = 3; predicted 7.80 mg/mL, 95% prediction interval 3.82–11.79 mg/mL). Its yield matched that of the 0.4 M centre points (13.28 mg/mL), whereas the four other 0.6 M S-2P runs gave 2.3–8.8 mg/mL and the eGFP salt-screening run at the same conditions gave 6.9 mg/mL. Run 17 was reproducible within the salt-screening campaign, so the discrepancy did not arise from a single replicate, but it was performed at an earlier stage of column use and outside the response-surface run sequence. It was therefore excluded from model fitting and is reported only as a salt-screening result (Figure 1D). Its exclusion removed the significant lack of fit of the fibre S-2P model (p = 0.001 with and 0.39 without run 17) and raised R² from 0.72 to 0.87 (Tables S3 and S4c). To show that the exclusion was not selective, the fibre models were also fitted with every run and with no salt-screening run (Table S4c): in all three fits the models were significant (p < 0.001), Gu-HCl accounted for 67–97% of the model sum of squares and the predicted Gu-HCl optimum lay at 0.35–0.40 M for both constructs. For each model, the tested run closest to the predicted optimum in coded units was identified as the near-optimal run (Table S5). For fibre S-2P without run 17, the predicted yield at 0.4 M Gu-HCl varied by less than 13% across the load-concentration and flow-rate ranges, and the run closest to the optimum (run 19) was a single unreplicated run; the replicated centre-point runs (n = 4), predicted at 98% of the maximum, were therefore retained as the near-optimal run (Table S5). The monolith, bead and membrane response-surface designs were run independently of the salt screen, so the inclusion of salt-screening runs, and the check described above, apply only to the fibre adsorber. Models were fitted and statistical tests performed in Python 3 with statsmodels 0.15 and SciPy 1.18.

**Table 2.** Factor levels (low, centre, high) of the face-centred central composite designs. Levels were the same for both constructs unless stated otherwise.

| **Factor** | **Monolith** | **Beads** | **Membrane** | **Fibre** |
|---|---|---|---|---|
| Gu-HCl [M] | 0.4, 0.6, 0.8 (eGFP);<br>0.2, 0.4, 0.6 (S-2P) | 0.4, 0.6, 0.8 | 0.2, 0.4, 0.6 | 0.2, 0.4, 0.6 |
| Load flow rate [mL/min] | 1, 3, 5 | 0.1, 0.3, 0.5 | 0.5, 1.0, 1.5 | 1.0, 1.5, 2.0 |
| Residence time [s] * | 60, 20, 12 | 600, 200, 120 | 24, 12, 8 | 24, 16, 12 |
| mRNA load conc. [mg/mL] | 0.2, 0.4, 0.6 | 0.2, 0.4, 0.6 | 0.1, 0.2, 0.3 | 0.1, 0.2, 0.3 |
| mRNA load amount [mg] eGFP/S-2P | 5/5 | 5/5 | 2/2 | 4.0-9.6/<br>2.2-8.0 |

\*Residence time is the bed volume divided by the load flow rate.

#### 2.4.3 Definitions and calculations

Recovered mass (yield) is the mRNA concentration in the eluate multiplied by the eluate volume and is reported per mL of bed (mg/mL). Recovery is the yield as a percentage of the mass loaded, and the load challenge is the mass loaded per mL of bed (mg/mL). Because breakthrough was observed only qualitatively in the flow-through UV signal and no breakthrough curves were recorded, the yield is not a dynamic binding capacity (DBC) but a lower bound on the binding capacity under the conditions used; where nearly all of the applied mRNA was recovered, the yield was limited by the load rather than by the matrix. Residence time was calculated as bed volume divided by volumetric flow rate. Productivity was calculated as the yield per mL of bed per minute of total cycle time, the sum of the chromatographic run time (equilibration, load, wash, elution and cleaning-in-place) and a fixed 5 min allowance for preparation. The amortised cost model is described in Section 3.4 and its sensitivity analysis in Section 3.5; a normalised relative-cost comparison of the same runs is given in Table S11. Unless stated otherwise, values are means ± standard deviation of n = 3 (at the near-optimal runs, n = 4 for the fibre adsorber with both constructs and for the beads with S-2P; Table S5). Yields of the near-optimal runs (Section 3.3; Table S5) were compared between formats separately for each construct by one-way ANOVA on log-transformed yield, which equalised the variances (Brown–Forsythe test), followed by Tukey’s honestly significant difference (HSD) test; differences are reported as ratios of geometric means with 95% simultaneous confidence intervals. Welch’s ANOVA with the Games–Howell test on untransformed yield was used as a sensitivity analysis, and equivalence of the beads and monolith was assessed by two one-sided t-tests (TOST) with a ±10% margin. Differences were considered significant at p < 0.05 (Table S6).

#### 2.4.4 Amortised cost model

The cost of purifying 1 mg of mRNA with each format was estimated as

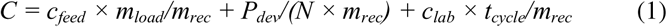

where *c_feed_* is the cost of the crude IVT mRNA applied to the column (USD/mg), *m_load_* and *m_rec_* are the masses loaded and recovered per cycle (Table 3), *P_dev_* is the device list price, *N* is the number of bind–elute cycles per device, *c_lab_* is a combined labour and facility rate (USD/h) and *t_cycle_* is the cycle time including the 5 min preparation allowance (Table 3). The three terms correspond to the raw-material, device and labour-and-facility categories. Baseline values were c_feed = USD 60/mg, representative of laboratory-scale IVT with research-grade reagents, and c_lab = USD 60/h, and the device list prices were USD 833 (monolith), 440 (beads), 500 (membrane) and 274 (fibre adsorber) (Table S10). The three cost inputs and the number of cycles per device were varied in a deterministic one-at-a-time sensitivity analysis, in which each input was set to a series of discrete levels while the other three were held at their baseline values (Section 3.5). No probability distribution was assigned to any input and no sampling scheme was used. The levels, ranges and their basis are given in Table S7 and the resulting costs in Table S8. Because Eq. (1) is additive in cfeed, clab and Pdev, the only pair of inputs that interacts is the device price with N; a variance decomposition over the same ranges confirmed that first-order Sobol indices account for essentially all of the variance (Table S9), so a sampled or variance-based design would be expected to reproduce the one-at-a-time result. Device list prices and their sources are listed in Table S10. mRNA that was not recovered was treated as lost. This is conservative for formats with lower recovery, because unbound mRNA in the flow-through can in principle be re-applied. Buffers, chromatography system and quality-control costs were assumed equal across formats and were excluded. The break-even number of cycles between two formats was obtained by equating Eq. (1) for both formats and solving for *N*. The model structure follows our earlier techno-economic analyses of RNA vaccine manufacturing [29,30].

**Table 3.** Productivity of the four matrices at the near-optimal runs (Table S5).

|  | **Monolith column [1 mL]** | **Beads column [1 mL]** | **Membrane column [0.2 mL]** | **Fibre column [0.4 mL]** | **mRNA construct** |
|---|---|---|---|---|---|
| Amount of mRNA recovered each run [mg]<sup>(A)</sup> | 3.700 | 3.960 | 1.392 | 7.170 | eGFP |
|  | 3.370 | 3.783 | 1.128 | 5.310 | S-2P |
| Amount of mRNA loaded each run [mg]<sup>(B)</sup> | 5.00 | 5.00 | 2 | 8.15 | eGFP |
|  | 5.00 | 5.00 | 2 | 7.20 | S-2P |
| Recovery of the applied load [%]<sup>(C)</sup> | 74.0 | 79.2 | 69.6 | 88.0 | eGFP |
|  | 67.4 | 75.7 | 56.4 | 73.8 | S-2P |
| Chromatography matrix volume [mL]<sup>(D)</sup> | 1 | 1 | 0.2 | 0.4 | eGFP/S-2P |
| Load challenge [mg mRNA/mL matrix]<sup>(E)</sup> | 5.0 | 5.0 | 10 | 20.4 | eGFP |
|  | 5.0 | 5.0 | 10 | 18.0 | S-2P |
| Time consumed for chromatography run [min]<sup>(F)</sup> | 28.3 | 109.3 | 26.4 | 52.1 | eGFP |
|  | 32.5 | 67.7 | 42.0 | 35.3 | S-2P |
| Preparation time [min]<sup>(G)</sup> | 5 | 5 | 5 | 5 | eGFP/S-2P |
| Productivity [mg mRNA/mL matrix/min] for each run<sup>(H)</sup> | 0.111 | 0.035 | 0.221 | 0.314 | eGFP |
|  | 0.090 | 0.052 | 0.120 | 0.329 | S-2P |

(A) The mass of mRNA recovered from the chromatography test with different columns, quantified by UV spectrometry.
(B) The amount of mRNA applied to the column in each run, calculated as load volume multiplied by load concentration, as given in Table 2.
C) Recovery, the ratio of mRNA recovered (A) to mRNA loaded (B). Loads were set in advance for each format and not adjusted to breakthrough. Where the load challenge (E) was close to or below the capacity of the bed, as for the fibre adsorber with eGFP, nearly all of the applied mRNA was captured and recovery is high; where it exceeded the capacity, recovery is bounded by the capacity of the matrix. Recovery is therefore not an independent measure of elution efficiency.
(D) The bed volume of the chromatography matrix applied in the experiment.
(E) The load challenge is the mass of mRNA loaded (B) divided by the matrix volume (D).
(F) The time consumed for the chromatography process, including start-up equilibration, load, wash, elution and shut-down column cleaning.
(G) The time consumed for preparing buffers, samples, equipment and consumables.
(H) The productivity is calculated as (A) divided by the product of the matrix volume (D) and the total cycle time, that is, the sum of the run time (F) and the preparation time (G).

**Table 4.** Cost per mg (USD) of different columns at N = 1, 20, and 100. Grouped by RNA construct.

|  | eGFP |  |  |  | S-2P |  |  |  |
|---|---|---|---|---|---|---|---|---|
| N | Monolith | Beads | Membrane | Fibre | Monolith | Beads | Membrane | Fibre |
| 1 | 315.1 | 215.7 | 467.7 | 114.4 | 347.5 | 214.9 | 591.3 | 140.5 |
| 20 | 101.3 | 110.1 | 126.4 | 78.1 | 112.7 | 104.4 | 170.2 | 91.5 |
| 100 | 92.3 | 105.7 | 112.1 | 76.5 | 102.8 | 99.8 | 152.5 | 89.4 |

## 3. Results

### 3.1 Recovered mRNA mass as a function of loading salt type and concentration

Cations screen the electrostatic repulsion between the phosphate backbones of the poly(A) tail and the oligo-dT ligand, and ionic strength is therefore a primary determinant of oligo-dT binding capacity [7,9], while chaotropic Gu-HCl additionally destabilises mRNA secondary structure and increases the accessibility of the tail [7]. Figure 1 compares the mass of mRNA recovered per mL of matrix from each format as a function of NaCl and Gu-HCl concentration in the loading buffer.

With NaCl (Figure 1A, B), every format showed an optimum in yield whose position depended on both the matrix and the construct. For eGFP the monolith and beads required 1.0 M NaCl to reach their maxima (3.6 and 3.9 mg/mL), whereas the membrane peaked at 0.4 M (6.3 mg/mL) and the fibre adsorber at 0.6 M (17.1 mg/mL). The longer S-2P construct required less salt: the monolith and beads peaked at 0.6 M (2.9 and 3.2 mg/mL) and the membrane and fibre adsorber at 0.4 M (6.4 and 9.3 mg/mL). Above these optima the yield fell on every format, but at very different rates. Taking one 0.2 M step past the optimum as a common measure, eGFP mass fell by 3% on the beads, 6% on the membrane, 20% on the monolith and 23% on the fibre adsorber, whereas the corresponding S-2P losses were 5%, 28%, 33% and 53%. The beads were therefore the most tolerant of excess salt for both constructs, holding 2.3-3.9 mg/mL of eGFP and 1.8-3.2 mg/mL of the S-2P across the whole 0.2-1.2 M range.

Gu-HCl increased the peak yield relative to NaCl in seven of the eight matrix–construct combinations and shifted the optimum to a lower concentration in five of them (monolith with both constructs, beads and fibre adsorber with eGFP, membrane with S-2P); the optimum was unchanged for the beads with S-2P, the membrane with eGFP (0.4 M in both salts) and the fibre adsorber with S-2P (Figure 1C, D). The monolith reached 4.05 mg eGFP/mL at 0.6 M and 3.78 mg S-2P/mL at 0.4 M; the beads 4.47 mg eGFP/mL at 0.8 M and 4. 03 mg S-2P/mL at 0.6 M; and the membrane 8.74 mg eGFP/mL at 0.4 M and 7.79 mg S-2P/mL at 0.2 M. The fibre adsorber again recovered the most mRNA per unit volume, 16.0 mg eGFP/mL and 15.0 mg S-2P/mL at 0.4 M, roughly twice the membrane and four times the monolith and beads. Above the optimum matrices diverged: the monolith and beads declined gradually and retained 2–3.5 mg/mL at 0.8–1.0 M; the membrane retained less than 2 mg/mL at 0.8 M for either construct; and the fibre adsorber lost more than half of its recovered eGFP mass between 0.4 and 0.6 M (16.0 to 6.9 ± 0.4 mg/mL) while retaining most of its S-2P yield at 0.6 M in the salt screen (12.9 ± 0.6 mg/mL). This S-2P value was not reproduced in the response-surface runs at 0.6 M (2.3–8.8 mg/mL), and a model fitted to those runs predicted 7.8 mg/mL at the same conditions, so it was excluded from the response-surface model (Section 2.4.2) and the S-2P yield of the fibre adsorber at 0.6 M Gu-HCl remains uncertain.

Two features distinguished the fibre adsorber. First, its optimum lay at a low-to-moderate salt concentration, 0.4 M Gu-HCl for both constructs and 0.4–0.6 M NaCl, similar to the membrane (0.2–0.4 M) and lower than the monolith and beads with eGFP (0.6–1.0 M), consistent with the manufacturer’s recommendation of moderate-salt loading [22]. Second, for the short eGFP construct NaCl was at least as effective as Gu-HCl (17.1 versus 16.0 mg/mL), whereas for every other matrix–construct combination, including the fibre adsorber with S-2P (9.3 versus 15.0 mg/mL), Gu-HCl was clearly superior.

### 3.2 Response-surface mapping of load flow rate, load concentration and Gu-HCl concentration

The effects of the three process parameters (PPs), Gu-HCl concentration, load flow rate and mRNA load concentration, were mapped for each matrix and construct with response-surface designs (Section 2.4; factor ranges in Table 2), with the Gu-HCl window set from the salt screen above. The quadratic models were significant for all eight matrix–construct pairs (p ≤ 0.014; Table S3). They described the monolith, the membrane with S-2P, the beads with S-2P and the fibre adsorber with both constructs well (R² 0.79–0.94, adjusted R² 0.74–0.92), whereas the beads with eGFP and the membrane with eGFP gave nearly flat surfaces relative to the replicate scatter (R² 0.41 and 0.47; predicted R² ≤ 0.11), so their optima are poorly defined. Lack of fit was significant for all models except those for the fibre adsorber (p = 0.63 for eGFP and 0.39 for S-2P), because the replicate scatter was small relative to the deviations from a quadratic surface; the models therefore describe the relative effects of the process parameters better than they predict absolute yields. The response surfaces should accordingly be read as interpretive rather than predictive, and the optima of the two weakest models, the beads with eGFP and the membrane with eGFP, should not be used to locate operating conditions. Gu-HCl was the dominant factor for the fibre adsorber (97% of the model sum of squares for eGFP and 84% for S-2P), the membrane with S-2P (78%), the monolith with S-2P (60%) and the beads with eGFP (57%) (Table S4b). For the fibre adsorber, the membrane with S-2P and the monolith with S-2P, a strongly negative quadratic Gu-HCl term made the yield rise to a construct-and matrix-specific optimum before falling away, whereas for the beads with eGFP the yield increased with Gu-HCl up to the upper limit of the range (0.8 M). In the other three combinations Gu-HCl accounted for only 21–36% of the model sum of squares: load concentration dominated for the membrane with eGFP (52%) and, through its curvature and its interaction with flow rate, for the beads with S-2P (63% together), and flow rate together with its interaction with Gu-HCl dominated for the monolith with eGFP (58%).

As shown in Figure 2, the model-predicted maxima within the design space (Table S3) were 3.8 mg eGFP/mL for the monolith (0.69 M Gu-HCl, 0.2 mg/mL, 4.4 mL/min) and 3.4 mg S-2P/mL (0.46 M, 0.27 mg/mL, 1.0 mL/min); 4.2 mg eGFP/mL for the beads (0.8 M, 0.6 mg/mL, 0.26 mL/min) and 3.7 mg S-2P/mL (0.61 M, 0.39 mg/mL, 0.3 mL/min); 7.8 mg eGFP/mL for the membrane (0.6 M, 0.3 mg/mL, 1.1 mL/min, at the upper edge of the Gu-HCl and load-concentration ranges and poorly defined, although the salt screen in Figure 1C gave higher recoveries at 0.4 M) and 6.4 mg S-2P/mL (0.37 M, 0.23 mg/mL, 0.5 mL/min); and 18.4 mg eGFP/mL for the fibre adsorber (0.35 M, 0.23 mg/mL, 1.0 mL/min) and 13.4 mg S-2P/mL (0.40 M, 0.13 mg/mL, 2.0 mL/min, at the edge of the load-concentration and flow-rate ranges and poorly defined, because the predicted S-2P yield at 0.4 M Gu-HCl varied only between 11.7 and 13.4 mg/mL across those ranges). The optimal Gu-HCl concentration was therefore highest for the beads (0.6–0.8 M), followed by the monolith (0.46–0.69 M), the membrane (0.37–0.6 M) and the fibre adsorber (0.35–0.40 M). Every run on the fibre column at 0.4 M Gu-HCl gave more than 15 mg eGFP/mL and, with one exception (9.9 mg/mL at 0.1 mg/mL and 1.0 mL/min), more than 11.8 mg S-2P/mL irrespective of the other two factors. Every fibre run at 0.6 M Gu-HCl was also accompanied by a rise in column pressure during loading (Section 4), so the descending arm of the fibre Gu-HCl response cannot be attributed to binding chemistry alone.

**Figure 2.**
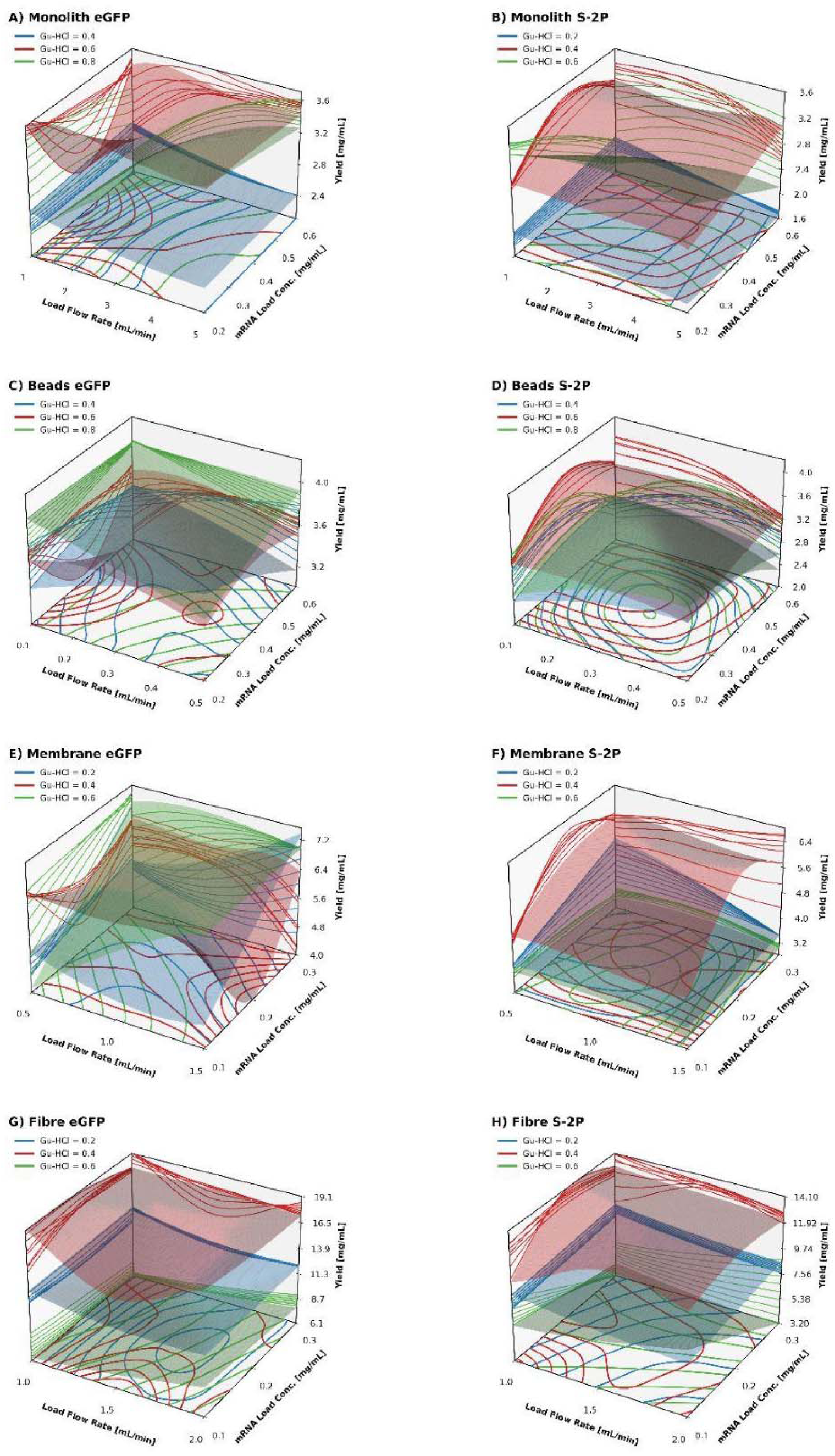
Response surfaces of the recovered mRNA mass as a function of load flow rate, mRNA load concentration and Gu-HCl concentration for the four oligo-dT matrices with eGFP (left) and S-2P (right) mRNA. Rows, top to bottom: monolith, beads, membrane and fibre adsorber (panels A–H). Each translucent surface is the fitted model (Table S4a) at one Gu-HCl concentration (colour-coded; ranges in Table 2); the fibre S-2P model (H) excludes run 17 (Section 2.4.2).

The response to flow rate separated the matrices by transport mechanism. The beads, operated at the lowest flow rates of the study (0.1–0.5 mL/min; residence times 600–120 s), gave their highest predicted yields at 0.26–0.3 mL/min (about 200 s), with a curved flow response (quadratic term p = 0.025 for eGFP), consistent with intra-particle transport limitations. The three convective formats were far less flow-sensitive, but not insensitive: the monolith showed no significant main effect of flow rate with S-2P (linear term p = 0.09, although the Gu-HCl × flow-rate interaction was significant, p = 0.018) but, with eGFP, gained yield with increasing flow rate at the higher Gu-HCl concentrations (Gu-HCl × flow-rate interaction, p < 0.001) across 1–5 mL/min (residence times 60–12 s), whereas the membrane favoured low-to-moderate flow rates (0.5–1.1 mL/min; 24–11 s) and the fibre adsorber gained little with eGFP and modestly with S-2P on halving the flow rate from 2.0 to 1.0 mL/min (12 to 24 s; 17.7 to 17.9 mg eGFP/mL and 12.7 to 14.0 mg S-2P/mL), with a significant Gu-HCl × flow-rate interaction for eGFP (p = 0.011), although for S-2P the difference lay within the replicate scatter and flow rate accounted for less than 0.1% of the model sum of squares. Load concentration had little effect on the fibre adsorber, the beads and the monolith with eGFP (0.1%, 4% and 4% of the model sum of squares) and a moderate effect on the monolith with S-2P (14%, with a significant Gu-HCl × load-concentration interaction, p < 0.001). It was, however, the largest single factor for the membrane with eGFP, whose yield rose with load concentration (linear term p = 0.001), and it produced a pronounced optimum near 0.4 mg/mL for the beads with S-2P (quadratic term p < 0.001). On the fibre adsorber with S-2P, load concentration accounted for 3% of the model sum of squares (linear and quadratic terms p ≥ 0.09), but its interaction with Gu-HCl was significant (p < 0.001), with lower load concentrations favoured at higher Gu-HCl concentrations.

### 3.3 Comparison at near-optimal conditions

For each format and construct, the tested response-surface run closest to the model-predicted optimum in coded units was taken as the near-optimal run (Table S5; for the fibre adsorber with S-2P, the replicated centre-point runs, Section 2.4.2), and these runs are compared in Figure 3. They were performed at 0.6 M Gu-HCl, 0.2 mg/mL and 3 mL/min (eGFP) and 0.4 M, 0.4 mg/mL and 1 mL/min (S-2P) on the monolith; 0.8 M, 0.6 mg/mL and 0.1 mL/min (eGFP) and 0.6 M, 0.4 mg/mL and 0.3 mL/min (S-2P) on the beads; 0.6 M, 0.3 mg/mL and 1.5 mL/min (eGFP) and 0.4 M, 0.2 mg/mL and 0.5 mL/min (S-2P) on the membrane; and 0.4 M, 0.2 mg/mL and 1.0 mL/min (eGFP) and 0.4 M, 0.2 mg/mL and 1.5 mL/min (S-2P) on the fibre adsorber, with the load mass of each format as in Table 2. At these settings the models predicted 94–100% of the maximum, and the observed yields were 88–101% of the predicted maxima (Table S5). The monolith recovered 3.70 ± 0.21 mg eGFP/mL and 3.37 ± 0.25 mg S-2P/mL, the beads 3.96 ± 0.11 and 3.78 ± 0.24 mg/mL, the membrane 6.96 ± 0.48 and 5.64 ± 0.43 mg/mL and the fibre adsorber 17.9 ± 1.7 and 13.3 ± 2.4 mg/mL, so eGFP yields exceeded S-2P yields on every format. Yield differed between formats for both constructs (one-way ANOVA on log-transformed yield, p < 0.001; Table S6). The fibre adsorber gave 2.6-and 2.3-fold the yield of the membrane for eGFP and S-2P (95% CI 2.2–3.0 and 1.8–3.1), 4.5-and 3.5-fold that of the beads (3.8–5.3 and 2.7–4.5) and 4.8-and 3.9-fold that of the monolith (4.1–5.7 and 2.9–5.2; all p < 0.001; Tukey HSD on log yield). The membrane gave 1.8-and 1.5-fold the yield of the beads (1.5–2.1 and 1.1–2.0; p < 0.001 and p = 0.007) and 1.9-and 1.7-fold that of the monolith (1.6–2.2 and 1.2–2.3; p < 0.001 and p = 0.002). The beads and monolith did not differ significantly (ratios 1.07 and 1.12; 95% CI 0.90–1.27 and 0.85–1.49; p = 0.62 and 0.61), but their equivalence within ±10% could not be demonstrated (TOST, p = 0.24 and 0.65). Welch’s ANOVA with the Games–Howell test on untransformed yield gave the same pattern. The ranking of the formats by yield per mL of bed (fibre adsorber > membrane > beads and monolith, the last two not significantly different) was the same in the salt screening (Figure 1), the response surfaces (Figure 2) and the near-optimal runs (Figure 3), which indicates that it is robust to the operating conditions explored, with the caveat that the salt screen and the response-surface designs were not run at identical loads for the membrane and the fibre adsorber (Section 2.4.1). These ratios nevertheless compare yield at the applied loads, which differed between formats: the monolith and beads were challenged with only about a quarter of the fibre load (5.0 versus 20.4 mg/mL of bed for eGFP; Table 3), so the comparison does not establish the relative binding capacities, which would require breakthrough curves at a common residence time.

**Figure 3.**
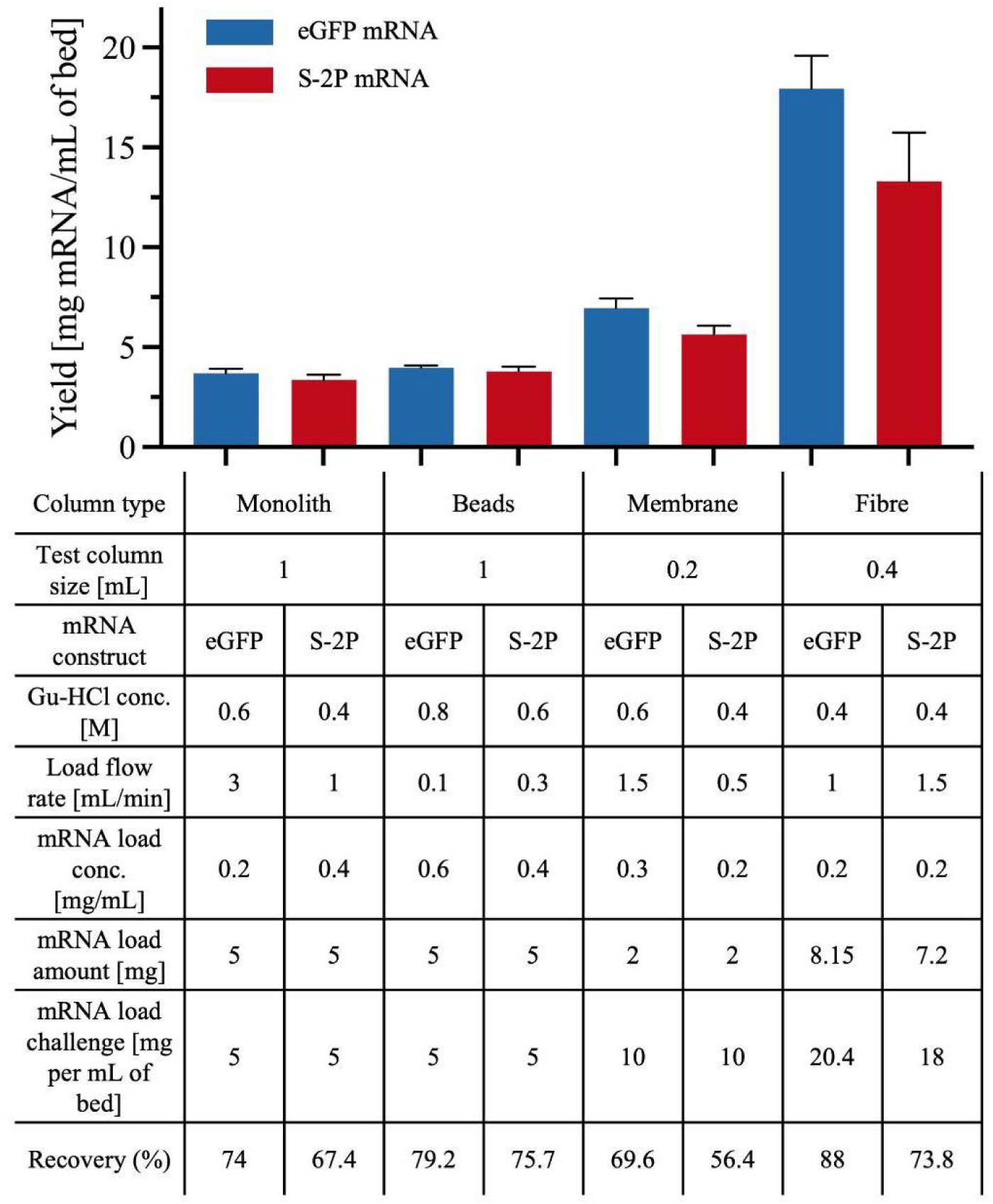
Recovered mass of eGFP (blue) and S-2P (red) mRNA on the four oligo-dT columns at the near-optimal response-surface runs (Table S5), whose process parameters are listed in the table beneath the chart together with the load applied, the load challenge and the recovery. Error bars show standard deviations (n = 3; n = 4 for the fibre adsorber with both constructs and for the beads with S-2P).

Productivity, defined as mRNA recovered per mL of bed per minute of total cycle time (chromatographic run plus 5 min preparation), is compared in Table 3 for the near-optimal runs and ranged about 9-fold across the four matrices for eGFP (0.314 to 0.035 mg/mL/min). The fibre adsorber recovered the most mRNA per cycle of any format, 7.17 mg eGFP and 5.31 mg S-2P from a 0.4 mL bed, that is, 1.8-and 1.4-fold the mass recovered by the 1 mL bead column (3.96 and 3.78 mg) and 1.9-and 1.6-fold that recovered by the 1 mL monolith (3.70 and 3.37 mg) from beds 2.5 times larger, and 5.2-and 4.7-fold the mass recovered by the 0.2 mL membrane (1.39 and 1.13 mg) from a bed half the size. Delivering 8.15 mg of eGFP at 0.2 mg/mL and 1.0 mL/min occupies 41 min on its own, so the fibre run takes 52 min and, with the 5 min preparation, the complete cycle 57 min; on that basis the fibre adsorber still gave the highest productivity of the study, 0.314 mg/mL/min for eGFP. For S-2P the 7.2 mg load, applied at 1.5 mL/min, takes 24 min to deliver, giving a 35 min run, a 40 min cycle and a productivity of 0.329 mg/mL/min. The membrane, whose small bed offsets its modest absolute recovery, was second at 0.221 and 0.120 mg/mL/min (26 and 42 min runs); the monolith reached 0.111 and 0.090 mg/mL/min (28 and 33 min); and the beads, whose low flow rates stretched the runs to 109 and 68 min, were lowest at 0.035 and 0.052 mg/mL/min. The fibre adsorber’s productivity was thus 1.4-and 2.7-fold that of the membrane, 2.8-and 3.7-fold that of the monolith and 9.1-and 6.3-fold that of the beads for eGFP and S-2P, respectively. For eGFP its advantage over the membrane is smaller than its advantage in yield per mL of bed (2.6-fold), because the large load needed to exploit that capacity dominates the cycle; for S-2P, whose membrane run was loaded at only 0.5 mL/min, it is larger (2.3-fold in yield). The monolith, beads and membrane runs exceed the time needed to deliver the load by 20, 26 and 22 min respectively (equilibration, wash, elution and cleaning), whereas the fibre runs exceed it by only about 11 min; this difference reflects the shorter step programme of the fibre adsorber, which is designed for rapid cycling: its supplier specifies a complete cycle (equilibration, load, wash, elution, strip, CIP and re-equilibration) of less than 15 min and recommends CIP only every one to five cycles [22]. Had the fibre adsorber required the same 22 min as the membrane, its productivity would have been about 0.26 mg/mL/min for both eGFP and S-2P, still 1.2-and 2.2-fold that of the membrane; this is the more conservative comparison of the two matrices.

### 3.4 Amortised cost per milligram of purified mRNA

Figure 4 shows the cost of purifying 1 mg of mRNA as a function of the number of cycles per device. For single use (N = 1), the device price dominated the differences between formats. The fibre adsorber was the least expensive (USD 114 and 140 per mg for eGFP and S-2P respectively), followed by the beads (216 and 215), the monolith (315 and 348) and the membrane (468 and 591) (**Figure 4A & B**). Because the device term falls as 1/N, these differences narrowed quickly with re-use. At N = 20, the fibre adsorber remained the least expensive (78 and 92 USD/mg), followed by the monolith (101 and 113) and the beads (110 and 104), while the membrane remained the most expensive (126 and 170 USD/mg) (Figure 5). The fibre adsorber was the least expensive format at every N for both constructs; the monolith became less expensive than the beads from about eight cycles for eGFP, whereas for S-2P the beads were less expensive than the monolith at every N.

**Figure 4.**
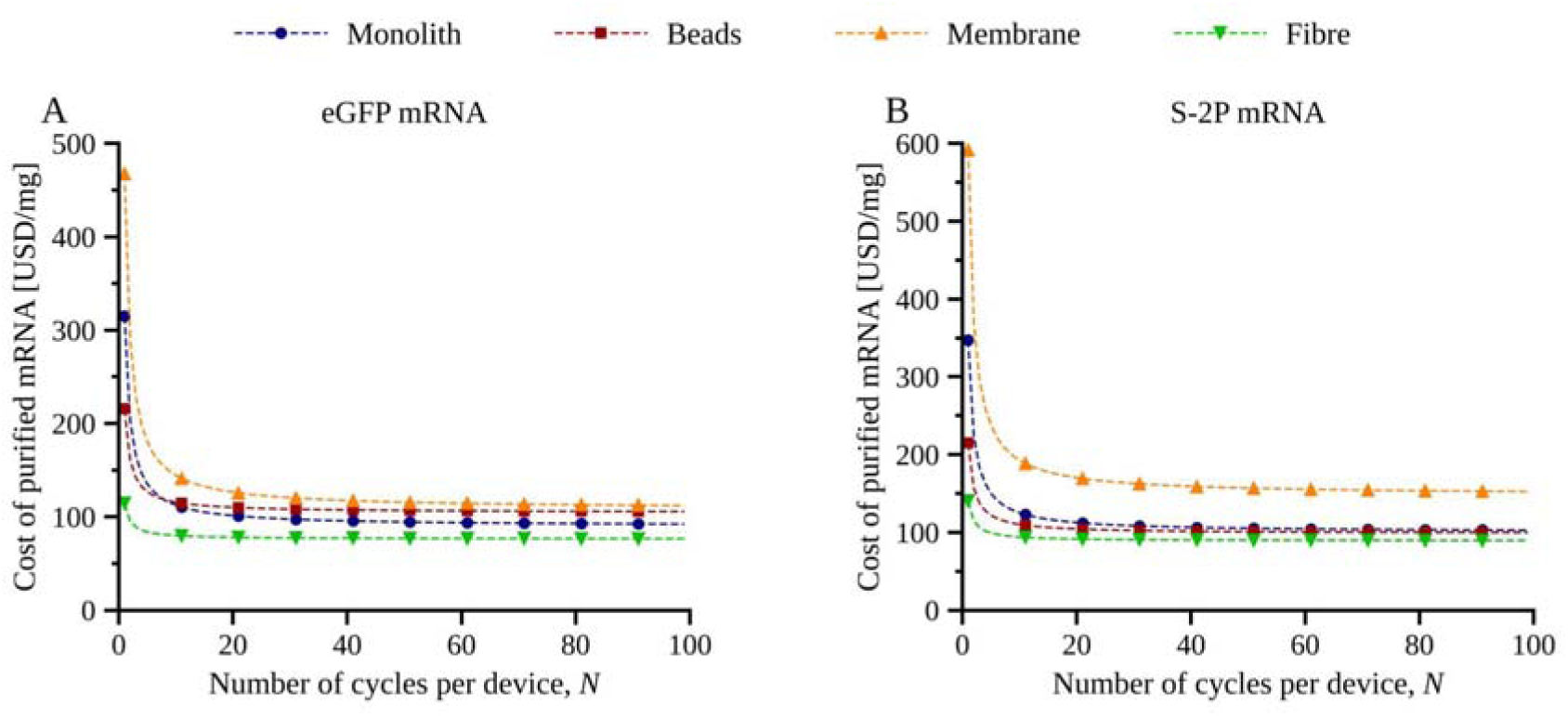
Amortised cost of purified mRNA (USD/mg) as a function of the number of cycles per device for the four column types with eGFP (A) and S-2P (B) mRNA, calculated with Eq. (1) (c_feed = USD 60/mg, c_lab = USD 60/h).

**Figure 5.**
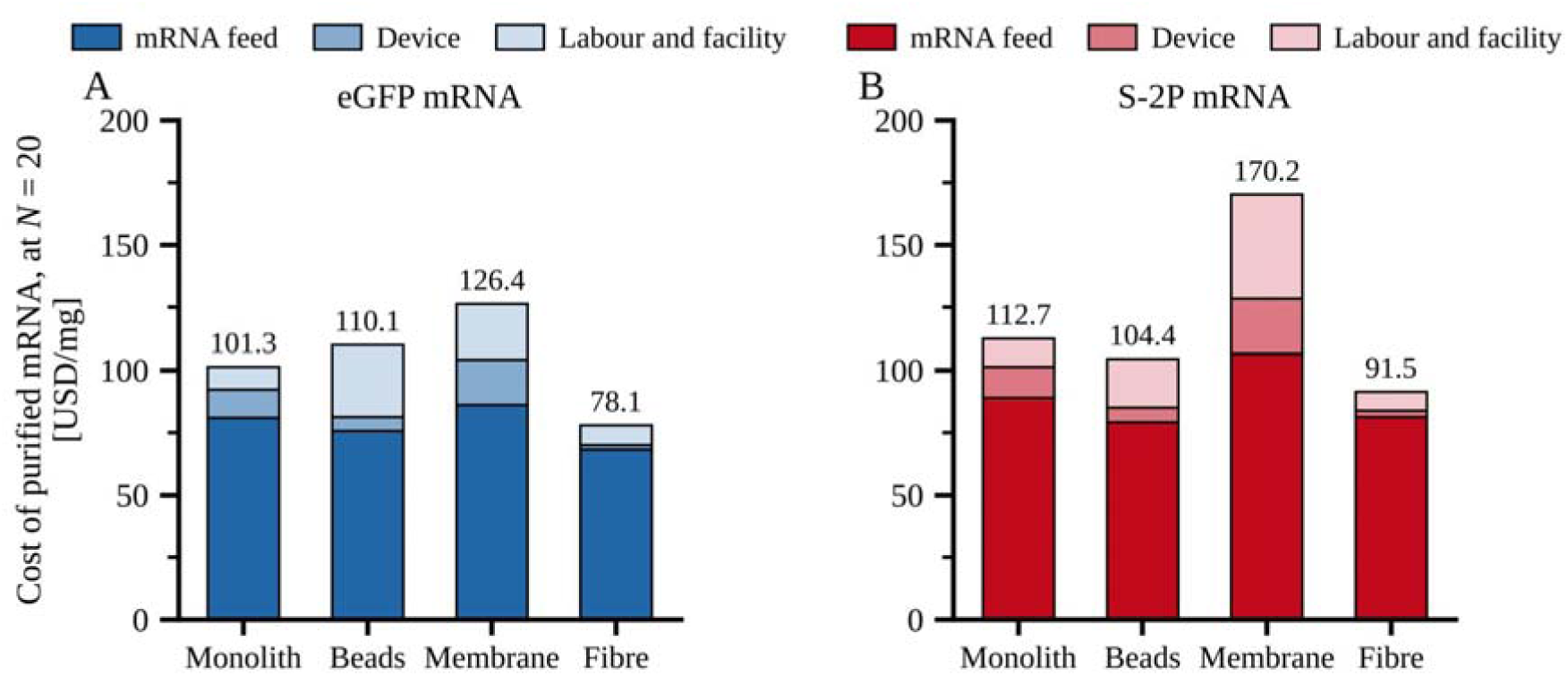
Breakdown of the cost per mg of purified mRNA (USD/mg) into mRNA feed, device, and labour and facility contributions with eGFP (A) and S-2P (B) mRNA, at N = 20 for eGFP and S-2P.

At N = 20, the mRNA feed accounted for 63–89% of the cost across formats (Figure 5). Beyond a few cycles, the ranking was therefore set mainly by recovery and by cycle time. The long cycle time of the beads raised their labour and facility cost to about USD 29 and 19 per mg for eGFP and S-2P, about 1.7 to 3.6 times that of the monolith (9 and 11) and the fibre adsorber (8 and 8). The membrane combined the lowest recovery (69.6 and 56.4%) with the highest device cost per milligram and, for S-2P, the highest labour and facility cost (about USD 42 per mg), because only 1.1–1.4 mg of mRNA was recovered per cycle from its 0.2 mL bed.

A normalised relative-cost comparison of the same near-optimal runs, which scores each format against the least efficient on raw-material use and against the monolith on device price, was also carried out and gives the same ranking at the baseline; because it combines categories normalised to different references and treats labour and facility cost as equal across formats, it is reported in Table S11 and Eq. (1) is used as the cost analysis throughout.

### 3.5 Sensitivity of the amortised cost model to its inputs

The three cost inputs of Eq. (1) and the number of cycles per device are estimates rather than measured quantities, so each was varied one at a time about the baseline while the others were held constant (Figure 6; levels and their basis in Table S7, full results in Table S8). The mRNA feed cost was by far the most influential input. Raising it from USD 6 to 120/mg changed the cost per mg of purified mRNA by 130 to 202 USD/mg, depending on the format. The labour and facility rate had the next largest effect: raising it from USD 15 to 120/h changed the cost per mg by 13 to 73 USD/mg. Changing the number of cycles per device from 10 to 400 changed it by 4 to 43 USD/mg. The device price mattered least: changing it by 50% in either direction changed the cost per mg by only 2 to 22 USD/mg. Figure S2 shows these swings for each input and each format. A full-factorial check was then run over the same four ranges, combining 58 feed costs, 8 labour and facility rates, 5 device prices and 6 cycle counts, which gives 13,920 combinations for each construct (Table S7). It found the fibre adsorber to be the least expensive format in every combination for eGFP and in 98.6% of combinations for S-2P, the exceptions requiring a feed cost of USD 90/mg or more together with the lowest labour and facility rate and 50 or more cycles, where the beads were less expensive by at most 1.1 USD/mg. Two orderings below the fibre adsorber are conditional. For eGFP the monolith and the beads exchange places at a labour and facility rate near USD 34/h at N = 20, the beads being less expensive below it, and the monolith overtakes the beads from 6 to 13 cycles depending on the feed cost. For S-2P the monolith overtakes the beads only beyond about 180 cycles at a feed cost of USD 45/mg, and not at all at USD 60/mg or above. The membrane was the most expensive format for S-2P in every combination tested; for eGFP it was the most expensive at the baseline, but the beads overtook it in 15% of combinations, all of which combine a feed cost at or below about USD 72/mg with 20 or more cycles per device, where the shrinking feed and device terms leave the long cycle time of the beads as the dominant cost. The device price, which separates the formats most clearly in a single-use comparison, accounted for 2.4 to 14.2% of the cost per mg at N = 20 and 0.1 to 0.8% at N = 400, so its influence falls away as the devices are re-used.

**Figure 6.**
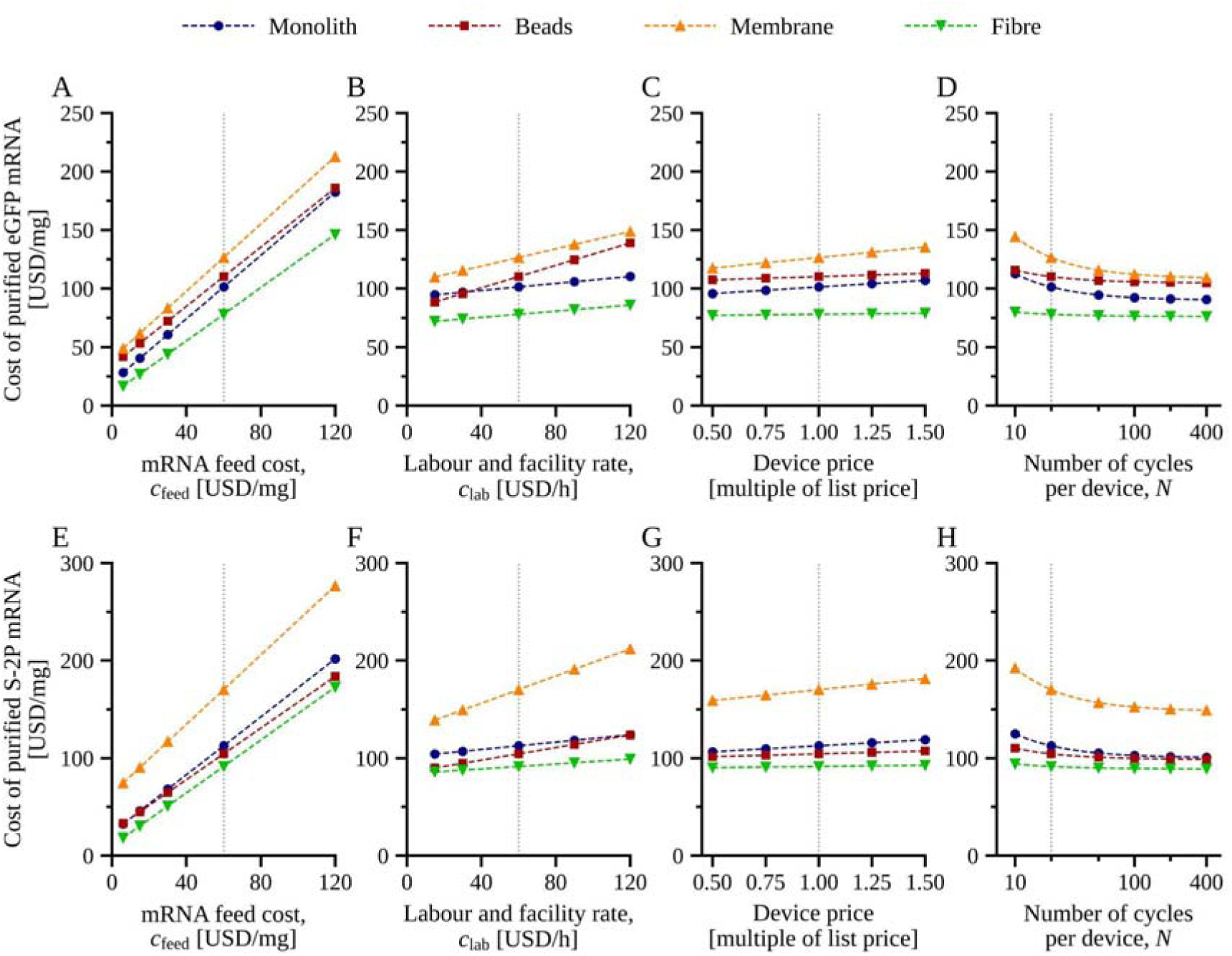
Sensitivity of the cost per mg of purified mRNA to the inputs of Eq. (1), for eGFP (A to D) and S-2P (E to H). Each panel varies one input over the range given in Table S7 while the other three are held at their baseline values (c_feed_ = USD 60/mg, c_lab_ = USD 60/h, device at supplier list price, N = 20 cycles per device), which the dotted vertical line marks. Panels A and E vary the mRNA feed cost, B and F the labour and facility rate, C and G the device price as a multiple of the supplier list price, and D and H the number of cycles per device on a logarithmic scale. The analysis is deterministic and one-at-a-time; no distribution is assigned to any input.

## 4. Discussion

In this study, we mapped the operating windows of four commercially available oligo-dT matrices for two mRNAs spanning the typical length range of vaccine and therapeutic constructs, and compared the matrices by yield, productivity and cost per milligram of purified mRNA.

The membrane and fibre adsorbers present their ligands on the surface of an open, predominantly convective network with a very high surface-area-to-volume ratio and short diffusion distances, so that a poly(A) tail encounters oligo-dT rapidly even at residence times of 10–25 s [19]. Electrospun cellulose fibres additionally lack dead-end pores and have a well-defined inter-fibre pore structure [19], which is consistent with a yield 2.3-to 2.6-fold that of the membrane at residence times of the same order (8–24 s) and, qualitatively, with the 2–4-fold and 2–13-fold advantages over the monolith and beads reported in a binding-capacity study of prototype fibre material [15]; the 3.5-to 4.8-fold advantage found here compares yield at unequal loads and is not directly comparable. The manufacturer quotes a dynamic binding capacity of at least 6 mg/mL for a 2000-nt mRNA in the recommended KCl loading buffer [22], whereas the yield of 13–18 mg/mL of bed obtained here, with different mRNAs and in Gu-HCl–phosphate buffer, indicates that the fibre bed can bind at least this much mRNA under these conditions, well above the quoted specification, and that loading-buffer optimisation is the key to exploiting it (the two values are not directly comparable because the DBC definition and the mRNA length differ). The POROS beads, although perfusive, still require mRNA to enter a pore network by combined convection and diffusion; their improvement at 0.1–0.3 mL/min (residence times of several minutes) is consistent with this transport limitation and with the strong residence-time dependence of beaded oligo-dT resins reported recently [16]. The monolith, with continuous macroporous channels, showed no significant main effect of flow rate with S-2P and, with eGFP, maintained or increased its yield up to 3–5 mL/min under elevated salt, as also observed by [15]; its plateau at high load concentration may point to ligand saturation rather than transport limitation [8]. Across the convective formats, the construct-size difference may also reflect an orientation-limited association step: as we have argued elsewhere, at short residence times the apparent on-rate for poly(A)–oligo-dT binding can be limited by the rotational and conformational diffusion required to present the comparatively small poly(A) segment to a surface-bound ligand, even where bulk mass transfer in the convective pore network is no longer limiting [31].

Ligand chemistry is likely to contribute significantly to the capacity of oligo-dT columns. The beads and membrane carry 25-mer oligo-dT, the monolith an 18-mer, and the fibre adsorber a proprietary oligo-dT (prototyped as a 20-mer) attached at a single 5′ point at more than 6 µmol/g fibre [15,22]. The two model constructs differ not only in total transcript length (930 vs 4284 nt) but also in poly(A) tail length (49 nt for eGFP and 120 nt for S-2P). Because the poly(A) tail is the primary hybridisation handle for oligo-dT capture, tail length directly affects duplex thermodynamic stability and the number of oligo-dT ligands each molecule can simultaneously engage [9,12]. Construct-dependent effects observed in this study, such as S-2P consistently reaching its binding optimum at lower Gu-HCl concentrations than eGFP across most formats, therefore cannot be attributed to transcript length alone; the more stable duplex formed by the 2.4-fold longer poly(A) tail of S-2P is an equally plausible explanation, consistent with the nearest-neighbour thermodynamic arguments developed below. A fully paired 25-mer duplex forms about 50 hydrogen bonds compared with 36 for an 18-mer; hydrogen bonding contributes only a few kJ/mol per base pair, but base stacking, counter-ion condensation and configurational entropy add to the net binding free energy [9,10]. Because poly(A)–oligo-dT is an RNA/DNA hybrid, RNA/DNA nearest-neighbour parameters are the appropriate reference [11,12]: these place the free energy of the rAA/dTT nearest-neighbour step (5′-rApA-3′/3′-dTpT-5′) at −0.7 kcal/mol (about −2.9 kJ/mol) at 37 °C in 100 mM NaCl (ΔH° = −7.8 kcal/mol, ΔS° = −22.9 cal mol−1 K−1; Banerjee et al., 2020), which corresponds to about −1.0 kcal/mol (−4.1 kJ/mol) at 25°C, close to the temperature used here, and at about −1.0 kcal/mol at 37 °C in 1 M NaCl (Sugimoto et al., 1995). The seven additional rA·dT steps of a 25-mer relative to an 18-mer would therefore add roughly 20–28 kJ/mol of stabilisation in 100 mM NaCl. This is an order-of-magnitude estimate that neglects the effects of Gu-HCl, of surface immobilisation and of the spacer. Longer ligands should therefore raise the affinity of each ligand, but capacity is set jointly by ligand density and accessibility, matrix geometry, residence time and buffer composition [8]. The monolith carries a moderate, well-accessed ligand load (about 0.08 µmol/mL) in convective macropores; the beads can be more heavily functionalised (about 0.3 µmol/mL) but kinetic limits to pore access give them a capacity similar to the monolith; and the membrane offers several times the specific surface area of the monolith (about 30,000 versus 5,000 and 1,000 cm²/g for membrane, monolith and beads) but only twice its capacity, consistent with the lower grafting density needed to preserve permeability at high flux [25–27]. The fibre adsorber combines high specific surface area with high ligand density, which is a plausible explanation for its higher yield per mL of bed than the membrane at similar residence times, although its ligand density, given per gram, cannot be compared directly with the per-mL values of the other formats. Expressed on a molar basis, the fibre adsorber recovered about 60 nmol of eGFP mRNA and 10 nmol of S-2P mRNA per mL of bed (taking 321.5 g/mol per nucleotide), a 6.2-fold difference for a 1.4-fold difference in mass. Before attributing this difference solely to accessible surface area or steric footprint, two alternative explanations merit consideration. First, the poly(A) tail length difference (49 nt for eGFP versus 120 nt for S-2P) means that each S-2P molecule can simultaneously engage more oligo-dT ligands than each eGFP molecule: a 120-nt tail can in principle hybridise with up to six 18–20-mer ligand chains, whereas a 49-nt tail can engage only two or three. Because the S-2P poly(A) tail is a split tail, its contiguous stretches are shorter than 120 nt, so six is an upper bound on the number of ligands it could engage. On a per-molecule basis, each S-2P therefore occupies more ligand sites, so fewer S-2P molecules than eGFP molecules would be expected per unit bed volume at saturation, independently of any steric or surface-area argument. Second, it is worth considering whether a 49-nt poly(A) tail is already long enough to saturate the available ligands under these conditions: if binding is limited by the three-dimensional accessibility of the tail within the folded mRNA rather than by the number of adenosine residues available, then extending tail length beyond a certain threshold would confer no additional benefit, and the stoichiometric advantage of S-2P may not fully materialise in practice. Literature directly addressing the minimum poly(A) tail length required to saturate oligo-dT affinity matrices is limited, and this question warrants further investigation. Taken together with the observation that the fibre adsorber recoveries at the applied loads (88.0% for eGFP and 73.8% for S-2P) indicate that the eGFP load approached and the S-2P load exceeded the binding capacity, the argument for a steric or surface-area limit is strongest if these loads were near breakthrough; without explicit breakthrough curves, the relative contributions of ligand stoichiometry, tail accessibility and steric footprint cannot be fully separated.

Gu-HCl gave a higher peak recovery than NaCl, and did so at equivalent molar concentrations, in seven of the eight matrix–construct combinations, consistent with its action on both of the barriers to hybridisation. As a monovalent salt it screens the electrostatic repulsion between the phosphate backbones of the poly(A) tail and the immobilised oligo-dT. As a chaotrope, Gu-HCl additionally lowers the stability of intramolecular mRNA secondary structure and so increases the fraction of poly(A) tails that are exposed and able to hybridise, whereas increasing NaCl tends to stabilise such structures [7]. All four matrices showed a capacity optimum followed by a decline at higher salt. On the beads and monolith the fall above 0.6–0.8 M Gu-HCl was gradual and is consistent with Gu-HCl beginning to weaken the A–T duplex itself and with salt-dependent compaction of the mRNA. On the membrane and fibre adsorber the loss was abrupt (from 8.7 to 1.9 mg/mL between 0.4 and 0.8 M on the membrane with eGFP, from 16.0 to 6.9 mg/mL between 0.4 and 0.6 M on the fibre adsorber with eGFP) and, on the fibre adsorber, every 0.6 M run showed a pressure rise during loading. One possible mechanism is salt-induced condensation of mRNA in the presence of spermidine, a trivalent polyamine present at millimolar concentrations in the IVT reaction mixture as a standard cofactor for T7 RNA polymerase and carried over into the diluted load at sub-millimolar concentrations. Multivalent cations such as spermidine can neutralise the phosphate backbone charge and condense or aggregate nucleic acids, although competing monovalent salt raises the polyamine concentration required for aggregation [32–34]; visible turbidity or precipitation in crude IVT loads at higher salt concentrations is consistent with this mechanism and has been observed in our hands. Aggregated mRNA or spermidine–mRNA co-precipitates could accumulate at the inlet of small-pore matrices (0.7 µm for the fibre bed; 0.45 µm for the membrane), increasing hydraulic resistance while leaving bound mRNA in place, consistent with the simultaneous pressure rise and capacity loss observed above 0.4 M Gu-HCl [35]. The monolith and beads, with their larger and more rigid pore networks, would be less vulnerable to blockage of this kind. If this mechanism is operative, in-line 0.2-µm prefiltration of the load before application to thin-bed formats would be a straightforward mitigation, removing aggregates before they reach the matrix and aligning with the broader recommendation for tighter feed-quality control in GMP operation discussed below.

Productivity ranked fibre > membrane > monolith > beads, driven by the yield per unit volume and by the flow rate each matrix tolerates. For eGFP, the margin between the fibre adsorber and the membrane is nevertheless narrower than capacity alone would suggest, because the high capacity of the fibre bed can only be realised by applying a correspondingly large load, and at 0.2 mg/mL and 1.0 mL/min the loading step alone occupies about 41 min of the 57 min cycle (52 min run plus 5 min preparation). Capacity converts into productivity only to the extent that the feed can be delivered fast enough to use it, which for the fibre adsorber would require either a more concentrated feed or a higher load flow rate than the one that maximised its yield. The response-surface analysis indicates only a modest penalty in yield on doubling the load flow rate from 1.0 to 2.0 mL/min for the fibre adsorber (about 2% for eGFP and 9% for S-2P; Section 3.2), so a higher load flow rate may raise productivity substantially: loading the same mass at 2.0 mL/min and 0.3 mg/mL would shorten the load step from 41 to about 14 min and give a productivity of about 0.5–0.6 mg/mL/min, depending on how much of the yield is maintained. These productivity estimates might change in a GMP compliant environment where preparation could take 30-60 minutes instead of the 5 minutes accounted for here.

The membrane’s productivity was offset by raw-material losses (recoveries of 69.6% and 56.4% for eGFP and S-2P), which gave it the highest raw-material cost of the four formats. The losses may arise from unbound mRNA in the flow-through and early wash at high flux, but no flow-through, wash or strip fractions were analysed, so this remains a hypothesis to be tested by a mass balance of the near optimal runs; effects over repeated cycles, such as fouling, were not examined. Capacity, productivity and cost determine which format is technically and economically preferable, but the format actually deployed in a clinical or commercial process must also be available at scale, manufactured under an appropriate quality system and supported by the documentation that regulators expect for a chromatography material [36]. On these criteria the four tested matrices are at very different stages of maturity, and the ranking is almost the reverse of the ranking by yield.

The monolith and the beads are established manufacturing formats. CIMmultus Oligo-dT columns are offered from 1 mL to 8 L (40 L on request), with the 1 mL to 8 L sizes manufactured as cGMP-compliant columns supplied with integrity-test procedures, a defined sanitisation and CIP regime (0.5 M NaOH) and a demonstrated lifetime of at least 40 cleaning cycles at pressures up to 1.8–2 MPa [37,38]. POROS Oligo (dT)25 is supplied as bulk resin from 10 mL to 10 L for packing into standard bioprocess columns, tolerates 10 MPa, and pack sizes of 1 L and above are accompanied by a regulatory support package covering change control, lot release and resin qualification for IND and BLA submissions [27]. Both materials are stated by their suppliers to be suitable for GMP manufacturing [27,38] and so a process developed on them at the 1 mL scale used here could in principle be transferred to production scale by conventional linear scale-up (constant bed height and residence time for the beads; constant column-volume-normalised flow for the radial-flow monolith) without a change of material, subject to verification at scale.

The membrane and fibre adsorbers, by contrast, were evaluated here as research-grade devices. The Purexa OdT Maxi column is a laboratory-scale unit [25]; its manufacturer announced a manufacturing-grade Purexa line produced in an ISO 9001, ISO Class 7 clean-room facility in July 2025, with the anion-exchange chemistry released first and the remaining chemistries, including OdT, and the pilot-and process-scale devices to follow during 2025–2026 [39]. The Fibro dT adsorber is currently supplied only as the 0.4 mL unit used in this study, intended for research and process development; GMP-compliant formats for clinical manufacturing are stated to be under development [40]. The underlying fibre platform has been scaled for protein A from 0.4 mL to 2.4 L units with predictable performance across scales [41–43], and 0.4 mL Fibro dT units used here showed no measurable loss of yield over 118 bind-elute cycles (Figure S3), which indicates that scale-up and re-use of the fibre matrix are technically feasible, but neither the ligand-specific devices nor the accompanying validation package exist yet for the oligo-dT chemistry. Neither thin-bed format therefore offers today a materially identical, quality-system-manufactured device at the scale of a clinical batch, and any process developed on them would have to be re-qualified when such devices become available.

Four further aspects distinguish the matrices in a GMP setting. First, scale-up geometry: the monolith and beads scale by bed volume within a single housing, whereas membranes and fibre beds scale by frontal area and by stacking or parallelising units, so that flow distribution, hold-up volume and inter-unit variability become validation items in their own right [20]. Second, cleaning and re-use versus single use: the monolith and beads are designed for validated multi-cycle re-use with NaOH sanitisation, which suits high-value products where resin cost is amortised over many batches, whereas the membrane and fibre devices are conceived as single-use or limited-use consumables, which removes cleaning validation and carry-over risk but requires extractables and leachable data and a secure supply chain for every batch [44]. Third, robustness: the pressure rise and capacity loss observed on the fibre adsorber above 0.4 M Gu-HCl illustrate that thin-bed formats have a narrower window of operation with respect to feed quality and salt concentration, so that in-line clarification and tighter control of the load would need to be built into a GMP process, whereas the monolith and beads lost yield only gradually above their salt optima; in-line 0.2-µm prefiltration of the load is therefore specifically recommended for the thin bed matrices when Gu-HCl loading buffers are used in GMP operation. Fourth, process chemistry: all optimised conditions in this study use Gu-HCl in the loading buffer. Guanidinium is a potent protein denaturant and is classified as a process-related impurity requiring clearance to acceptable limits before formulation [45,46]; residual Gu-HCl in the eluate, which is collected as a water fraction, would need to be quantified and shown to fall below regulatory limits, and a dedicated clearance study would be required for any GMP process. NaCl, by contrast, is a pharmacopoeial reagent with no intrinsic toxicity concern, and where the yield at the NaCl optimum is within an acceptable range, it represents a simpler regulatory path. This trade-off between chromatographic performance and process-chemistry simplicity should be factored into buffer selection for clinical manufacture.

The fibre adsorber had nearly the same optimal conditions for both constructs tested (0.4 M Gu-HCl and 0.2 mg/mL load concentration, with 1.0 mL/min for eGFP and 1.5 mL/min for S-2P; model-predicted Gu-HCl optima 0.35–0.40 M, with the S-2P optimum poorly defined in load concentration and flow rate), suggesting that its operating window is relatively insensitive to construct identity, at least across the mRNA size and poly(A) tail length range examined here. This platform-like behaviour could reduce the process development burden when multiple mRNA sequences are to be purified on the same equipment, indicating the highest level of platformability out of these four OdT column formats, although more investigation and evaluation is needed. Moreover, the fibre column, due to its rapid cycling, appears most amenable to continuous multi-column processing.

This study has several limitations. First, the quality of the eluates (integrity, dsRNA and residual template DNA) was not assessed, so the comparison addresses capture performance rather than impurity clearance. Second, the loads were fixed in advance for the monolith, beads and membrane but set by breakthrough or by column blockage at higher chaotrope concentrations, for the fibre adsorber, so the yields are not binding capacities measured on a common basis. Third, one unit of each format was characterised at the 0.2–1 mL scale, and performance over repeated cycles was not examined systematically. Fourth, the cost analysis relies on public list prices and on the assumption that every format sustains the same number of cycles, which is the one input the sensitivity analysis cannot use to separate the formats because varying it moves all four together. These points define the future work set out in Section 5.

## 5. Conclusion

Four commercially available oligo-dT affinity matrices, monolith, perfusive beads, membrane and electrospun fibre, were characterised with a common experimental approach for a short and a long mRNA. Four conclusions emerge. First, the fibre adsorber gave the highest yield per mL of bed (17.9 mg eGFP and 13.3 mg S-2P per mL, 2.3-to 4.8-fold the other formats) and the highest productivity (0.31 and 0.33 mg/mL/min for eGFP and S-2P), followed by the membrane, monolith and beads, although these yields were obtained at load challenges that differed between formats (5–20 mg/mL of bed) and are not binding capacities. Second, Gu-HCl gave higher yields than NaCl in seven of the eight matrix–construct combinations, but the thin-bed membrane and fibre formats lost yield sharply above 0.4–0.6 M Gu-HCl, which sets an upper salt limit for their operation, whereas the monolith and beads declined gradually. Third, the fibre adsorber was the least expensive format per milligram of purified mRNA at the baseline cost inputs, both for single use and at any number of re-use cycles, and the membrane the most expensive; these rankings held across almost the whole range of cost inputs examined (Section 3.5); with re-use, the mRNA feed became the largest cost item, so recovery became the main driver of cost. Fourth, manufacturing readiness ranks the formats almost in reverse: only the monolith and beads are currently available as cGMP-compliant, process-scale materials with regulatory support, so the higher yield and productivity of the thin-bed formats can be exploited in clinical manufacturing only once manufacturing-grade devices and validation packages exist. Future work should measure eluate quality (integrity, dsRNA and residual DNA) and Gu-HCl clearance, extend lifetime studies to all four formats of columns, and verify the operating windows at pilot and GMP scale.

## Authorship contribution statement

**Jixin Qu**: Conceptualisation, Investigation, Methodology, Formal analysis, Writing – original draft, Writing – review & editing. **Manoj Pohare**: Conceptualisation, Investigation, Methodology, Formal analysis, Writing – original draft, Writing – review & editing. Jixin Qu and Manoj Pohare contributed equally to this work. **Shady Hamed**: Formal analysis, Validation, Visualization, Writing – review & editing. **Adithya Nair**: Investigation, Validation. **Mabrouka Maamra**: Project administration and resources. **Zoltán Kis**: Conceptualisation, Methodology, Supervision, Validation, Writing – review & editing, Funding acquisition.

## Acknowledgement

The authors thank Donaldson Company, Inc. (Purilogics), Cytiva, Sartorius BIA Separations and Thermo Fisher Scientific for providing the Purexa™ OdT, Fibro dT, CIMmultus® Oligo-dT and POROS™ GoPure Oligo (dT)25 columns used in this study free of charge or at substantially discounted prices. The suppliers had no role in the study design, data collection, analysis or interpretation, or the writing of the manuscript. The authors thank Kate A. Loveday for kindly providing a portion of the crude (unpurified) IVT reaction mixture used for the oligo-dT purification experiments in this study.

## Data availability

The data supporting this study, and the analysis code that reproduces the cost model and its sensitivity analysis, are available in the University of Sheffield research data repository (ORDA) at https://doi.org/10.15131/shef.data.35010028 [reserved DOI]. The deposit includes the raw replicate yields for every design run, the workbook behind the cost model, and a documented Python script that regenerates Tables S7 to S9 and Figure S2. A second script refits the fibre response-surface models and regenerates the fibre rows of Tables S3 to S5 and Table S4c. The record will be made public on acceptance of this article.

Until the record is made public on acceptance, reviewers can access it through this private link: https://figshare.com/s/186a11fc7323ee480eb2

## Declaration of generative AI in the manuscript preparation process

During the preparation of this work, the authors used Claude AI (Anthropic) to assist with the statistical and cost analyses, improve the language of the manuscript, and check reported values against the deposited data. The authors reviewed and verified all outputs and take full responsibility for the content of the published article.

