## Supplementary material for "Monolith, porous bead, membrane or fibre? A comparative characterisation of four oligo-dT affinity chromatography matrices for mRNA capture": Supplymentary information

**Supplementary information**

**Table S1.** Chromatography phases

| **Chromatography phases** | **Monolith column** | **Beads column** | **Membrane column** | **Fibre column** |
| --- | --- | --- | --- | --- |
| Equilibration (mL/min; CV) | 4;20 | 1;8 | 0.5; 15 | 3; 20 |
| Load (mL/min; CV) | Variable | Variable | Variable | Variable |
| Wash (mL/min; CV) | 4;20 | 1;8 | 0.5; 15 | 3; 15 |
| Elution (mL/min; CV) | 2;10 | 1;5 | 0.5; 10 | 3; 30 |
| CIP (mL/min; CV) | 2;10 | 1;5 | 0.5; 10 | 3; 20 |
| NaOH concentration for CIP (M) | 0.5 | 0.1 | 0.1 | 0.1 |
| Total non-load time (min) | 20.0 | 26.0 | 22.0 | 11.3 |

**Table S2.** Gu-HCl and NaCl salt screening conditions

| **Parameter** | **Monolith column** | **Beads column** | **Membrane column** | **Fibre column** | **RNA** |
| --- | --- | --- | --- | --- | --- |
| Load flow rate (mL/min) | 3 | 0.3 | 1.0 | 1.6 | eGFP |
|  | 3 | 0.3 | 1.0 | 1.6 | S-2P |
| Load Concentration (mg/mL) | 0.4 | 0.4 | 0.2 | 0.2 | eGFP |
|  | 0.4 | 0.4 | 0.2 | 0.2 | S-2P |
| Load mass (mg) | 5 | 5 | 2.5 | Varied upon NaCl/GuHCl concentration used in the experiments | eGFP |
|  | 5 | 5 | 2.5 |  | S-2P |

**Table S3.** Fit statistics of the quadratic response-surface models (yield, mg mRNA per mL of bed). "Settings" is the number of distinct factor-level combinations in the design, which is one fewer than the number of runs because the centre point is duplicated: 15 settings for the 16-run designs and 19 for the 20-run fibre designs. For fibre S-2P, salt-screening run 17 was excluded from the fit (Section 2.4.2 of the main text and Table S4c), leaving 36 observations at 18 settings.

| **Model** | **No. of obs.** | **Settings** | **R²** | **Adj. R²** | **Pred. R²** | **RMSE (mg/**  **mL)** | **Model F** | **Model p** | **LOF F** | **LOF df** | **LOF p** | **R² (run means)** | **Opt. Gu-HCl (M)** | **Opt. load conc. (mg/mL)** | **Opt. flow (mL/min)** | **Pred. max (mg/mL)** | **95% CI** | **Max. observed run mean (mg/mL)** |
| --- | --- | --- | --- | --- | --- | --- | --- | --- | --- | --- | --- | --- | --- | --- | --- | --- | --- | --- |
| Fibre – eGFP | 42 | 19 | 0.936 | 0.919 | 0.899 | 1.27 | 52.3 | <0.001 | 0.79 | 9, 23 | 0.63 | 0.98 | 0.35 | 0.23 | 1 | 18.41 | 17.40–19.42 | 19.18 |
| Fibre – S-2P | 36 | 18 | 0.870 | 0.825 | 0.780 | 1.50 | 19.4 | <0.001 | 1.14 | 8, 18 | 0.39 | 0.924 | 0.40 | 0.13 | 2.0 | 13.40 | 12.04–14.77 | 14.04 |
| Monolith – eGFP | 46 | 15 | 0.888 | 0.86 | 0.815 | 0.202 | 31.6 | <0.001 | 3 | 5, 31 | 0.025 | 0.96 | 0.69 | 0.2 | 4.4 | 3.83 | 3.66–4.00 | 3.7 |
| Monolith – S-2P | 46 | 15 | 0.844 | 0.805 | 0.75 | 0.29 | 21.7 | <0.001 | 10.09 | 5, 31 | <0.001 | 0.897 | 0.46 | 0.27 | 1 | 3.36 | 3.12–3.60 | 3.41 |
| Beads – eGFP | 46 | 15 | 0.411 | 0.264 | 0.013 | 0.281 | 2.8 | 0.014 | 4.43 | 5, 31 | 0.004 | 0.627 | 0.8 | 0.6 | 0.26 | 4.17 | 3.89–4.45 | 3.98 |
| Beads – S-2P | 46 | 15 | 0.79 | 0.738 | 0.666 | 0.257 | 15.1 | <0.001 | 6.52 | 5, 31 | <0.001 | 0.874 | 0.61 | 0.39 | 0.3 | 3.74 | 3.58–3.89 | 3.86 |
| Membrane – eGFP | 46 | 15 | 0.473 | 0.342 | 0.107 | 0.898 | 3.6 | 0.003 | 41.63 | 5, 31 | <0.001 | 0.509 | 0.6 | 0.3 | 1.1 | 7.77 | 6.89–8.66 | 7.36 |
| Membrane – S-2P | 46 | 15 | 0.877 | 0.846 | 0.804 | 0.484 | 28.5 | <0.001 | 11.97 | 5, 31 | <0.001 | 0.912 | 0.37 | 0.23 | 0.5 | 6.44 | 6.03–6.85 | 6.56 |

**Table S4a.** Coded (standardised) coefficients, significance and share of model sum of squares per term

| **Model** | **Term** | **Factor group** | **Coded coef. (mg/mL)** | **SE** | **t** | **p** | **Type III SS** | **Share of model SS** |
| --- | --- | --- | --- | --- | --- | --- | --- | --- |
| Fibre – eGFP | Gu-HCl | Gu-HCl | **-2.62** | 0.265 | -9.87 | **<0.001** | 157.1061 | 21.0% |
| Fibre – eGFP | Load conc. | Load conc. | -0.031 | 0.259 | -0.12 | 0.905 | 0.0234 | 0.0% |
| Fibre – eGFP | Flow rate | Flow rate | 0.079 | 0.252 | 0.31 | 0.757 | 0.1573 | 0.0% |
| Fibre – eGFP | Gu-HCl × Load conc. | Interactions | -0.231 | 0.317 | -0.73 | 0.471 | 0.8556 | 0.1% |
| Fibre – eGFP | Gu-HCl × Flow rate | Interactions | **0.853** | 0.317 | 2.69 | **0.011** | 11.6452 | 1.6% |
| Fibre – eGFP | Load conc. × Flow rate | Interactions | -0.136 | 0.284 | -0.48 | 0.634 | 0.3713 | 0.0% |
| Fibre – eGFP | Gu-HCl² | Gu-HCl | **-7.701** | 0.411 | -18.75 | **<0.001** | 566.4676 | 75.9% |
| Fibre – eGFP | Load conc.² | Load conc. | -0.325 | 0.474 | -0.68 | 0.499 | 0.7552 | 0.1% |
| Fibre – eGFP | Flow rate² | Flow rate | **1.116** | 0.472 | 2.37 | **0.024** | 9.0288 | 1.2% |
| Fibre – S-2P | Gu-HCl | Gu-HCl | **-1.663** | 0.364 | -4.58 | **<0.001** | 47.3619 | 14.6% |
| Fibre – S-2P | Load conc. | Load conc. | -0.535 | 0.302 | -1.77 | 0.088 | 7.1135 | 2.2% |
| Fibre – S-2P | Flow rate | Flow rate | 0.016 | 0.307 | 0.05 | 0.960 | 0.0059 | 0.0% |
| Fibre – S-2P | Gu-HCl × Load conc. | Interactions | **-1.408** | 0.376 | -3.74 | **<0.001** | 31.7110 | 9.8% |
| Fibre – S-2P | Gu-HCl × Flow rate | Interactions | 0.727 | 0.376 | 1.93 | 0.064 | 8.4463 | 2.6% |
| Fibre – S-2P | Load conc. × Flow rate | Interactions | -0.296 | 0.336 | -0.88 | 0.386 | 1.7553 | 0.5% |
| Fibre – S-2P | Gu-HCl² | Gu-HCl | **-5.715** | 0.573 | -9.98 | **<0.001** | 225.3828 | 69.6% |
| Fibre – S-2P | Load conc.² | Load conc. | -0.583 | 0.624 | -0.93 | 0.359 | 1.9741 | 0.6% |
| Fibre – S-2P | Flow rate² | Flow rate | 0.013 | 0.624 | 0.02 | 0.984 | 0.0009 | 0.0% |
| Monolith – eGFP | Gu-HCl | Gu-HCl | **0.19** | 0.037 | 5.15 | **<0.001** | 1.083 | 10.5% |
| Monolith – eGFP | Load conc. | Load conc. | -0.068 | 0.037 | -1.84 | 0.074 | 0.1387 | 1.3% |
| Monolith – eGFP | Flow rate | Flow rate | **0.258** | 0.037 | 6.99 | **<0.001** | 1.9969 | 19.4% |
| Monolith – eGFP | Gu-HCl × Load conc. | Interactions | -0.082 | 0.041 | -2 | 0.053 | 0.1633 | 1.6% |
| Monolith – eGFP | Gu-HCl × Flow rate | Interactions | **0.367** | 0.041 | 8.9 | **<0.001** | 3.2414 | 31.4% |
| Monolith – eGFP | Load conc. × Flow rate | Interactions | -0.027 | 0.041 | -0.67 | 0.51 | 0.0181 | 0.2% |
| Monolith – eGFP | Gu-HCl² | Gu-HCl | **-0.577** | 0.072 | -7.97 | **<0.001** | 2.596 | 25.2% |
| Monolith – eGFP | Load conc.² | Load conc. | **0.193** | 0.072 | 2.66 | **0.012** | 0.2893 | 2.8% |
| Monolith – eGFP | Flow rate² | Flow rate | **-0.317** | 0.072 | -4.38 | **<0.001** | 0.7842 | 7.6% |
| Monolith – S-2P | Gu-HCl | Gu-HCl | **0.363** | 0.053 | 6.86 | **<0.001** | 3.9531 | 29.7% |
| Monolith – S-2P | Load conc. | Load conc. | **-0.211** | 0.053 | -3.99 | **<0.001** | 1.3356 | 10.0% |
| Monolith – S-2P | Flow rate | Flow rate | -0.093 | 0.053 | -1.76 | 0.087 | 0.2595 | 2.0% |
| Monolith – S-2P | Gu-HCl × Load conc. | Interactions | **-0.336** | 0.059 | -5.69 | **<0.001** | 2.7135 | 20.4% |
| Monolith – S-2P | Gu-HCl × Flow rate | Interactions | **0.146** | 0.059 | 2.47 | **0.018** | 0.5133 | 3.9% |
| Monolith – S-2P | Load conc. × Flow rate | Interactions | 0.006 | 0.059 | 0.11 | 0.916 | 0.0009 | 0.0% |
| Monolith – S-2P | Gu-HCl² | Gu-HCl | **-0.722** | 0.104 | -6.96 | **<0.001** | 4.0593 | 30.5% |
| Monolith – S-2P | Load conc.² | Load conc. | **-0.242** | 0.104 | -2.33 | **0.025** | 0.4558 | 3.4% |
| Monolith – S-2P | Flow rate² | Flow rate | -0.002 | 0.104 | -0.02 | 0.986 | 0 | 0.0% |
| Beads – eGFP | Gu-HCl | Gu-HCl | **0.184** | 0.051 | 3.58 | **<0.001** | 1.0157 | 47.3% |
| Beads – eGFP | Load conc. | Load conc. | -0.004 | 0.051 | -0.08 | 0.938 | 0.0005 | 0.0% |
| Beads – eGFP | Flow rate | Flow rate | 0.02 | 0.051 | 0.39 | 0.699 | 0.012 | 0.6% |
| Beads – eGFP | Gu-HCl × Load conc. | Interactions | 0.095 | 0.057 | 1.66 | 0.107 | 0.2166 | 10.1% |
| Beads – eGFP | Gu-HCl × Flow rate | Interactions | -0.065 | 0.057 | -1.13 | 0.265 | 0.1014 | 4.7% |
| Beads – eGFP | Load conc. × Flow rate | Interactions | -0.055 | 0.057 | -0.96 | 0.344 | 0.0726 | 3.4% |
| Beads – eGFP | Gu-HCl² | Gu-HCl | 0.165 | 0.101 | 1.63 | 0.111 | 0.2112 | 9.8% |
| Beads – eGFP | Load conc.² | Load conc. | 0.105 | 0.101 | 1.04 | 0.306 | 0.0853 | 4.0% |
| Beads – eGFP | Flow rate² | Flow rate | **-0.235** | 0.101 | -2.34 | **0.025** | 0.4314 | 20.1% |
| Beads – S-2P | Gu-HCl | Gu-HCl | 0.033 | 0.047 | 0.7 | 0.487 | 0.0327 | 0.7% |
| Beads – S-2P | Load conc. | Load conc. | -0.035 | 0.047 | -0.74 | 0.461 | 0.0368 | 0.8% |
| Beads – S-2P | Flow rate | Flow rate | -0.002 | 0.047 | -0.04 | 0.966 | 0.0001 | 0.0% |
| Beads – S-2P | Gu-HCl × Load conc. | Interactions | -0.074 | 0.053 | -1.4 | 0.169 | 0.1305 | 2.7% |
| Beads – S-2P | Gu-HCl × Flow rate | Interactions | -0.021 | 0.053 | -0.4 | 0.688 | 0.0108 | 0.2% |
| Beads – S-2P | Load conc. × Flow rate | Interactions | **-0.244** | 0.053 | -4.64 | **<0.001** | 1.4259 | 29.5% |
| Beads – S-2P | Gu-HCl² | Gu-HCl | **-0.408** | 0.092 | -4.43 | **<0.001** | 1.2978 | 26.9% |
| Beads – S-2P | Load conc.² | Load conc. | **-0.458** | 0.092 | -4.97 | **<0.001** | 1.6352 | 33.8% |
| Beads – S-2P | Flow rate² | Flow rate | -0.183 | 0.092 | -1.99 | 0.055 | 0.2614 | 5.4% |
| Membrane – eGFP | Gu-HCl | Gu-HCl | **0.432** | 0.164 | 2.64 | **0.012** | 5.5987 | 19.8% |
| Membrane – eGFP | Load conc. | Load conc. | **0.572** | 0.164 | 3.49 | **0.001** | 9.8155 | 34.7% |
| Membrane – eGFP | Flow rate | Flow rate | 0.252 | 0.164 | 1.54 | 0.133 | 1.9051 | 6.7% |
| Membrane – eGFP | Gu-HCl × Load conc. | Interactions | -0.015 | 0.183 | -0.08 | 0.935 | 0.0054 | 0.0% |
| Membrane – eGFP | Gu-HCl × Flow rate | Interactions | 0.165 | 0.183 | 0.9 | 0.374 | 0.6534 | 2.3% |
| Membrane – eGFP | Load conc. × Flow rate | Interactions | -0.065 | 0.183 | -0.35 | 0.725 | 0.1014 | 0.4% |
| Membrane – eGFP | Gu-HCl² | Gu-HCl | 0.176 | 0.322 | 0.55 | 0.587 | 0.2424 | 0.9% |
| Membrane – eGFP | Load conc.² | Load conc. | **0.796** | 0.322 | 2.48 | **0.018** | 4.9407 | 17.5% |
| Membrane – eGFP | Flow rate² | Flow rate | **-0.804** | 0.322 | -2.5 | **0.017** | 5.03 | 17.8% |
| Membrane – S-2P | Gu-HCl | Gu-HCl | **-0.356** | 0.088 | -4.03 | **<0.001** | 3.8021 | 8.8% |
| Membrane – S-2P | Load conc. | Load conc. | **0.236** | 0.088 | 2.67 | **0.011** | 1.6709 | 3.9% |
| Membrane – S-2P | Flow rate | Flow rate | -0.144 | 0.088 | -1.63 | 0.112 | 0.6221 | 1.4% |
| Membrane – S-2P | Gu-HCl × Load conc. | Interactions | **-0.235** | 0.099 | -2.38 | **0.023** | 1.3254 | 3.1% |
| Membrane – S-2P | Gu-HCl × Flow rate | Interactions | 0.185 | 0.099 | 1.87 | 0.069 | 0.8214 | 1.9% |
| Membrane – S-2P | Load conc. × Flow rate | Interactions | -0.165 | 0.099 | -1.67 | 0.104 | 0.6534 | 1.5% |
| Membrane – S-2P | Gu-HCl² | Gu-HCl | **-1.967** | 0.174 | -11.34 | **<0.001** | 30.1452 | 69.5% |
| Membrane – S-2P | Load conc.² | Load conc. | **-0.687** | 0.174 | -3.96 | **<0.001** | 3.6788 | 8.5% |
| Membrane – S-2P | Flow rate² | Flow rate | 0.293 | 0.174 | 1.69 | 0.1 | 0.6677 | 1.5% |

*Bold: p < 0.05.*

**Table S4b.** Share of model sum of squares by factor (linear + quadratic terms) and for all interactions

| **Model** | **Gu-HCl (%)** | **Load conc. (%)** | **Flow rate (%)** | **Interactions (%)** | **Check (sum)** |
| --- | --- | --- | --- | --- | --- |
| Fibre – eGFP | 96.9% | 0.1% | 1.2% | 1.7% | 100.0% |
| Fibre – S-2P | 84.2% | 2.8% | 0.0% | 12.9% | 100.0% |
| Monolith – eGFP | 35.7% | 4.2% | 27.0% | 33.2% | 100.0% |
| Monolith – S-2P | 60.3% | 13.5% | 2.0% | 24.3% | 100.0% |
| Beads – eGFP | 57.2% | 4.0% | 20.7% | 18.2% | 100.0% |
| Beads – S-2P | 27.5% | 34.6% | 5.4% | 32.4% | 100.0% |
| Membrane – eGFP | 20.6% | 52.2% | 24.5% | 2.7% | 100.0% |
| Membrane – S-2P | 78.2% | 12.3% | 3.0% | 6.5% | 100.0% |

**Sensitivity of the fibre response-surface models to run 17 and to the salt-screening runs**

Four salt-screening runs per construct, performed at 0.2, 0.4 and 0.6 M Gu-HCl with 0.2 mg/mL load concentration and 1.5 mL/min, coincided with design points of the fibre response-surface designs and were incorporated into the fibre datasets rather than repeated (eGFP runs 5, 10, 11 and 16; S-2P runs 5, 11, 12 and 17). The fibre models were fitted to three datasets: A, every run; B, every run except S-2P run 17, which is the analysis reported in the main text and in Tables S3 to S5; and C, the response-surface runs alone, without any salt-screening run. Each salt-screening run was then compared with the 95% prediction interval of the dataset C model for the mean of the same number of replicates, t(0.975, df) × √(SE(fit)² + s²/n), where SE(fit) is the standard error of the fitted mean, s² the residual mean square and n the number of replicates of the run. Dataset C contains no centre point, because both centre-point runs of each construct came from the salt screen, so its lack-of-fit test rests on fewer degrees of freedom. The eGFP model is the same in datasets A and B.

Seven of the eight salt-screening runs lay within the prediction interval (Table S4c, part B). The exception, S-2P run 17 at 0.6 M Gu-HCl, gave 12.85 mg/mL against a prediction of 7.80 mg/mL; its yield matched the 0.4 M centre points rather than the other 0.6 M runs (2.3–8.8 mg/mL). In all three datasets the models were significant (p < 0.001), Gu-HCl accounted for 67–97% of the model sum of squares and the predicted Gu-HCl optimum lay at 0.35–0.40 M for both constructs (part A). Lack of fit was significant only for S-2P with run 17 included (dataset A). The predicted S-2P maximum ranged from 13.4 to 14.6 mg/mL, and its position in load concentration and flow rate differed between datasets, consistent with the flat S-2P response at 0.4 M Gu-HCl (Table S5, note). The analysis is reproduced by the script fibre_doe_run17_sensitivity.py in the data deposit.

**Table S4c.** Sensitivity of the fibre response-surface models to the salt-screening runs incorporated into the fibre datasets. Part A: fit statistics and predicted optimum for each dataset; LOF, lack of fit. Part B: comparison of each salt-screening run with the 95% prediction interval of the model fitted without any salt-screening run (dataset C).

**A. Fit statistics and predicted optimum**

| **Dataset** | **Model** | **No. of obs.** | **Settings** | **R²** | **Adj. R²** | **Pred. R²** | **RMSE (mg/mL)** | **Model p** | **LOF p** | **Opt. Gu-HCl (M)** | **Opt. load conc. (mg/mL)** | **Opt. flow (mL/min)** | **Pred. max (95% CI) (mg/mL)** | **Gu-HCl share of model SS (%)** |
| --- | --- | --- | --- | --- | --- | --- | --- | --- | --- | --- | --- | --- | --- | --- |
| A and B. All runs | Fibre – eGFP | 42 | 19 | 0.936 | 0.919 | 0.899 | 1.27 | <0.001 | 0.63 | 0.35 | 0.22 | 1.0 | 18.42 (17.41–19.43) | 96.9 |
| C. No salt-screening runs | Fibre – eGFP | 30 | 16 | 0.952 | 0.930 | 0.899 | 1.18 | <0.001 | 0.30 | 0.35 | 0.30 | 1.0 | 18.61 (17.12–20.09) | 94.3 |
| A. All runs | Fibre – S-2P | 39 | 19 | 0.723 | 0.637 | 0.535 | 2.14 | <0.001 | 0.001 | 0.39 | 0.19 | 1.5 | 14.36 (13.00–15.72) | 67.3 |
| B. Run 17 excluded (adopted) | Fibre – S-2P | 36 | 18 | 0.870 | 0.825 | 0.780 | 1.50 | <0.001 | 0.39 | 0.40 | 0.13 | 2.0 | 13.40 (12.04–14.77) | 84.2 |
| C. No salt-screening runs | Fibre – S-2P | 29 | 16 | 0.905 | 0.860 | 0.785 | 1.38 | <0.001 | 0.25 | 0.36 | 0.19 | 1.5 | 14.58 (12.15–17.01) | 68.8 |

**B. Salt-screening runs against the dataset C model**

| **Model** | **Run** | **Gu-HCl (M)** | **Load conc. (mg/mL)** | **Flow (mL/min)** | **n** | **Observed yield, mean ± SD (mg/mL)** | **Predicted, dataset C model (mg/mL)** | **95% prediction interval for the mean of n (mg/mL)** | **Within interval** |
| --- | --- | --- | --- | --- | --- | --- | --- | --- | --- |
| Fibre – eGFP | 5 | 0.2 | 0.2 | 1.5 | 4 | 11.74 ± 1.87 | 10.68 | 7.38–13.97 | Yes |
| Fibre – eGFP | 10 | 0.4 | 0.2 | 1.5 | 3 | 16.95 ± 1.65 | 16.04 | 13.57–18.51 | Yes |
| Fibre – eGFP | 11 | 0.4 | 0.2 | 1.5 | 2 | 17.62 ± 2.23 | 16.04 | 13.38–18.70 | Yes |
| Fibre – eGFP | 16 | 0.6 | 0.2 | 1.5 | 3 | 6.88 ± 0.43 | 5.28 | 1.90–8.65 | Yes |
| Fibre – S-2P | 5 | 0.2 | 0.2 | 1.5 | 3 | 8.38 ± 0.03 | 11.38 | 7.39–15.36 | Yes |
| Fibre – S-2P | 11 | 0.4 | 0.2 | 1.5 | 2 | 13.60 ± 0.74 | 14.40 | 11.24–17.56 | Yes |
| Fibre – S-2P | 12 | 0.4 | 0.2 | 1.5 | 2 | 12.95 ± 4.10 | 14.40 | 11.24–17.56 | Yes |
| **Fibre – S-2P** | **17** | **0.6** | **0.2** | **1.5** | **3** | **12.85 ± 0.59** | **7.80** | **3.82–11.79** | **No** |

*Bold: run outside the prediction interval.*

**Table S5.** Model-predicted optima and the near-optimal experimental runs (replicate yields)

| **Model** | **Model-predicted optimum** | | | | **Near-optimal experimental run** | | | | | **RNA Yield (mg/mL bed)** | | | | | | | | | |
| --- | --- | --- | --- | --- | --- | --- | --- | --- | --- | --- | --- | --- | --- | --- | --- | --- | --- | --- | --- |
|  | Gu-HCl (M) | Load conc. (mg/mL) | Flow (mL/min) | Pred. max (mg/mL) | Run(s) | Gu-HCl (M) | Load conc. (mg/mL) | Flow (mL/min) | Coded distance | Rep 1 | Rep 2 | Rep 3 | Rep 4 | Mean | SD | n | Predicted at run | Pred. at run / pred. max | Observed / pred. max |
| Fibre – eGFP | 0.35 | 0.225 | 1 | 18.41 | 8 | 0.4 | 0.2 | 1 | 0.35 | 17.95 | 20.25 | 17 | 16.5 | 17.93 | 1.66 | 4 | 18.01 | 98% | 97% |
| Fibre – S-2P | 0.40 | 0.128 | 2.0 | 13.40 | 11,12 | 0.4 | 0.2 | 1.5 | 1.23 | 14.125 | 13.075 | 15.85 | 10.05 | 13.28 | 2.44 | 4 | 13.08 | 98% | 99% |
| Monolith – eGFP | 0.69 | 0.2 | 4.4 | 3.83 | 14 | 0.6 | 0.2 | 3 | 0.83 | 3.68 | 3.5 | 3.92 |  | 3.70 | 0.21 | 3 | 3.66 | 96% | 97% |
| Monolith – S-2P | 0.46 | 0.27 | 1 | 3.36 | 1 | 0.4 | 0.4 | 1 | 0.72 | 3.65 | 3.29 | 3.17 |  | 3.37 | 0.25 | 3 | 3.26 | 97% | 100% |
| Beads – eGFP | 0.8 | 0.6 | 0.26 | 4.17 | 6 | 0.8 | 0.6 | 0.1 | 0.8 | 3.84 | 3.98 | 4.06 |  | 3.96 | 0.11 | 3 | 4.02 | 96% | 95% |
| Beads – S-2P | 0.61 | 0.39 | 0.3 | 3.74 | 11,16 | 0.6 | 0.4 | 0.3 | 0.07 | 4.07 | 3.48 | 3.82 | 3.76 | 3.78 | 0.24 | 4 | 3.73 | 100% | 101% |
| Membrane – eGFP | 0.6 | 0.3 | 1.1 | 7.77 | 7 | 0.6 | 0.3 | 1.5 | 0.8 | 6.5 | 7.45 | 6.93 |  | 6.96 | 0.48 | 3 | 7.28 | 94% | 90% |
| Membrane – S-2P | 0.37 | 0.23 | 0.5 | 6.44 | 15 | 0.4 | 0.2 | 0.5 | 0.34 | 5.94 | 5.145 | 5.835 |  | 5.64 | 0.43 | 3 | 6.33 | 98% | 88% |

**Note.** For fibre S-2P, the model fitted without run 17 places the optimum at the edge of the design (0.40 M Gu-HCl, 0.128 mg/mL, 2.0 mL/min), and the run closest to it is run 19 (0.4 M, 0.1 mg/mL, 2.0 mL/min; coded distance 0.28), a single unreplicated run (11.85 mg/mL). Across the load-concentration and flow-rate ranges at 0.4 M Gu-HCl the predicted yield varies only between 11.7 and 13.4 mg/mL, and the centre point is predicted at 13.08 mg/mL, 98% of the maximum and within its 95% confidence interval (12.04–14.77 mg/mL). The replicated centre-point runs 11 and 12 (n = 4) were therefore retained as the near-optimal run, so that the between-format comparison in Table S6 rests on replicated data. Using instead run 19 (n = 1; 11.85 mg/mL), run 8 (0.4 M, 0.2 mg/mL, 2.0 mL/min; n = 2; 12.74 ± 1.08 mg/mL) or run 10 (0.4 M, 0.1 mg/mL, 1.5 mL/min; n = 3; 13.01 ± 1.21 mg/mL) would lower the fibre S-2P near-optimal yield by at most 11% and would not change the ranking of the formats.

**Table S6.** Between-format comparison of yield at the near-optimal runs (replicate data)

A. Near-optimal run yields (mg/mL bed)

| **Construct** | **Format** | **Run(s)** | **Mean yield [mg/mL of bed]** | **SD [mg/mL of bed]** | **CV (%)** | **n (replicate runs)** |
| --- | --- | --- | --- | --- | --- | --- |
| eGFP | Monolith | 14 | 3.7 | 0.21 | 5.7 | 3 |
| eGFP | Beads | 6 | 3.96 | 0.11 | 2.8 | 3 |
| eGFP | Membrane | 7 | 6.96 | 0.48 | 6.8 | 3 |
| eGFP | Fibre | 8 | 17.93 | 1.66 | 9.3 | 4 |
| S-2P | Monolith | 1 | 3.37 | 0.25 | 7.4 | 3 |
| S-2P | Beads | 11,16 | 3.78 | 0.24 | 6.4 | 4 |
| S-2P | Membrane | 15 | 5.64 | 0.43 | 7.7 | 3 |
| S-2P | Fibre | 11,12 | 13.28 | 2.44 | 18.3 | 4 |

B. Variance homogeneity and omnibus tests

| **Construct** | **Test** | **Test statistic** | **df** | **p** |
| --- | --- | --- | --- | --- |
| eGFP | Brown–Forsythe (raw yield) | 2.27 |  | 0.15 |
| eGFP | Brown–Forsythe (log yield) | 0.72 |  | 0.565 |
| eGFP | One-way ANOVA (log yield) | 409.21 | 3, 9 | <0.001 |
| S-2P | Brown–Forsythe (raw yield) | 3.2 |  | 0.071 |
| S-2P | Brown–Forsythe (log yield) | 1.1 |  | 0.395 |
| S-2P | One-way ANOVA (log yield) | 96.65 | 3, 10 | <0.001 |
| eGFP | One-way ANOVA (raw yield) | 166.42 | 3, 9.0 | <0.001 |
| eGFP | Welch ANOVA (raw yield) | 102.48 | 3, 4.4 | <0.001 |
| S-2P | One-way ANOVA (raw yield) | 43.94 | 3, 10.0 | <0.001 |
| S-2P | Welch ANOVA (raw yield) | 32.22 | 3, 5.0 | 0.001 |

C. Pairwise comparisons – primary: Tukey HSD on log yield

| **Construct** | **Comparison** | **Ratio of geometric means** | **95% CI** | **p (Tukey, log)** | **Difference (mg/mL)** | **p (Tukey, raw)** | **p (Games–Howell, raw)** |
| --- | --- | --- | --- | --- | --- | --- | --- |
| eGFP | Beads / Monolith | 1.07 | 0.90–1.27 | **0.623** | 0.26 | 0.988 | 0.386 |
| eGFP | Membrane / Monolith | 1.88 | 1.58–2.24 | <0.001 | 3.26 | 0.013 | 0.007 |
| eGFP | Fibre / Monolith | 4.83 | 4.11–5.69 | <0.001 | 14.23 | <0.001 | 0.001 |
| eGFP | Membrane / Beads | 1.76 | 1.48–2.09 | <0.001 | 3 | 0.021 | 0.016 |
| eGFP | Fibre / Beads | 4.51 | 3.84–5.31 | <0.001 | 13.97 | <0.001 | 0.001 |
| eGFP | Fibre / Membrane | 2.57 | 2.19–3.03 | <0.001 | 10.97 | <0.001 | 0.001 |
| S-2P | Beads / Monolith | 1.12 | 0.85–1.49 | **0.613** | 0.41 | 0.978 | 0.258 |
| S-2P | Membrane / Monolith | 1.67 | 1.24–2.27 | 0.002 | 2.27 | 0.235 | 0.01 |
| S-2P | Fibre / Monolith | 3.89 | 2.93–5.17 | <0.001 | 9.9 | <0.001 | 0.011 |
| S-2P | Membrane / Beads | 1.49 | 1.12–1.98 | 0.007 | 1.86 | 0.332 | 0.021 |
| S-2P | Fibre / Beads | 3.47 | 2.67–4.51 | <0.001 | 9.49 | <0.001 | 0.013 |
| S-2P | Fibre / Membrane | 2.33 | 1.75–3.09 | <0.001 | 7.64 | <0.001 | 0.021 |

**D. Equivalence of beads and monolith (TOST, margin ±10% of monolith mean)**

| **Construct** | **Difference (mg/mL)** | **90% CI** | **Margin (mg/mL)** | **p (TOST)** | **Conclusion** |
| --- | --- | --- | --- | --- | --- |
| eGFP | 0.26 | -0.06 to 0.58 | 0.37 | 0.241 | Not equivalent within ±10% |
| S-2P | 0.41 | 0.02 to 0.80 | 0.34 | 0.646 | Not equivalent within ±10% |

**Table S7.** Inputs varied in the sensitivity analysis of the amortised cost model, Eq. (1). The analysis is deterministic and one factor at a time: each input is set to the levels listed while the other three are held at their baseline values. No probability distribution is assigned to any input and no sampling scheme is used, so there is no Latin hypercube, Sobol sequence or Monte Carlo component. Table S9 verifies that this design is adequate.

The ranges above were also used for the full-factorial check of the ranking of the formats reported in Section 3.5 of the main text. That check evaluated Eq. (1) at every combination of 58 mRNA feed costs (USD 6 to 120/mg in steps of USD 2/mg), 8 labour and facility rates (USD 15, 30, 45, 60, 75, 90, 105 and 120/h), 5 device prices (0.5, 0.75, 1.00, 1.25 and 1.50 times the list price) and 6 cycle counts (10, 20, 50, 100, 200 and 400 cycles), which gives 58 × 8 × 5 × 6 = 13,920 combinations for each construct.

| **Variable** | **Symbol** | **Unit** | **Baseline** | **Range** | **Levels tested** | **Basis and reference** |
| --- | --- | --- | --- | --- | --- | --- |
| mRNA feed cost | c_feed_ | USD per mg | 60 | 6 to 120 | 6, 15, 30, 60, 120 | Baseline represents laboratory-scale IVT with research-grade reagents. The lower end approximates the drug-substance cost at manufacturing scale [[1,2]](https://www.zotero.org/google-docs/?TzheJD); the upper end doubles the baseline. |
| Labour and facility rate | c_lab_ | USD per h | 60 | 15 to 120 | 15, 30, 60, 90, 120 | Combined labour and facility rate. The lower end represents manufacturing in low- and middle-income countries; the upper end approximates a GMP cleanroom rate in a high-income setting [[1,2]](https://www.zotero.org/google-docs/?Uv8Dzh). |
| Device list price | P_dev_ | multiple of list price | 1.0 | 0.5 to 1.5 | 0.5, 0.75, 1.0, 1.25, 1.5 | Supplier list prices as of 27 October 2026 at a EUR/USD rate of 1.11 where applicable (Table S10). The range covers volume discounts and movement in list prices. |
| Cycles per device | N | bind-elute cycles | 20 | 10 to 400 | 10, 20, 50, 100, 200, 400 | At least 40 cleaning cycles are demonstrated for the monolith [[3,4]](https://www.zotero.org/google-docs/?vpmIb8); 0.4 mL units of the protein A version of the fibre platform, rather than the dT adsorber, retained a high dynamic binding capacity over 300 cleaning cycles [[5]](https://www.zotero.org/google-docs/?EEi0gA); 118 bind-elute cycles were run on the fibre unit used here (Figure S3). |

**Table S8.** Cost per mg of purified mRNA (USD/mg) at every level of the one-at-a-time sensitivity analysis defined in Table S7. Baseline rows are shown in bold. The fibre adsorber is the least expensive and the membrane the most expensive format in every row, so only the order of the monolith and the beads changes; the last column gives the cheaper of those two.

| **Construct** | **Symbol and unit of the input varied** | **Value of the input varied** | **Monolith - cost of purified mRNA [USD/mg]** | **Beads - cost of purified mRNA [USD/mg]** | **Membrane - cost of purified mRNA [USD/mg]** | **Fibre - cost of purified mRNA [USD/mg]** | **Cheaper of monolith and beads** | **Baseline** |
| --- | --- | --- | --- | --- | --- | --- | --- | --- |
| eGFP | c_feed_ [USD/mg] | 6 | 28.3 | 41.9 | 48.9 | 16.7 | Monolith |  |
| eGFP | c_feed_ [USD/mg] | 15 | 40.4 | 53.3 | 61.8 | 26.9 | Monolith |  |
| eGFP | c_feed_ [USD/mg] | 30 | 60.7 | 72.2 | 83.3 | 44.0 | Monolith |  |
| **eGFP** | **c_feed_ [USD/mg]** | **60** | **101.3** | **110.1** | **126.4** | **78.1** | **Monolith** | **yes** |
| eGFP | c_feed_ [USD/mg] | 120 | 182.3 | 185.9 | 212.6 | 146.3 | Monolith |  |
| eGFP | c_lab_ [USD/h] | 15 | 94.6 | 88.5 | 109.7 | 72.1 | Beads |  |
| eGFP | c_lab_ [USD/h] | 30 | 96.8 | 95.7 | 115.3 | 74.1 | Beads |  |
| **eGFP** | **c_lab_ [USD/h]** | **60** | **101.3** | **110.1** | **126.4** | **78.1** | **Monolith** | **yes** |
| eGFP | c_lab_ [USD/h] | 90 | 105.7 | 124.5 | 137.6 | 82.0 | Monolith |  |
| eGFP | c_lab_ [USD/h] | 120 | 110.2 | 138.9 | 148.7 | 86.0 | Monolith |  |
| eGFP | Device price | 0.5 x | 95.6 | 107.3 | 117.5 | 77.1 | Monolith |  |
| eGFP | Device price | 0.75 x | 98.4 | 108.7 | 121.9 | 77.6 | Monolith |  |
| **eGFP** | **Device price** | **1 x** | **101.3** | **110.1** | **126.4** | **78.1** | **Monolith** | **yes** |
| eGFP | Device price | 1.25 x | 104.1 | 111.5 | 130.9 | 78.5 | Monolith |  |
| eGFP | Device price | 1.5 x | 106.9 | 112.9 | 135.4 | 79.0 | Monolith |  |
| eGFP | N [cycles] | 10 | 112.5 | 115.7 | 144.4 | 80.0 | Monolith |  |
| **eGFP** | **N [cycles]** | **20** | **101.3** | **110.1** | **126.4** | **78.1** | **Monolith** | **yes** |
| eGFP | N [cycles] | 50 | 94.5 | 106.8 | 115.7 | 76.9 | Monolith |  |
| eGFP | N [cycles] | 100 | 92.3 | 105.7 | 112.1 | 76.5 | Monolith |  |
| eGFP | N [cycles] | 200 | 91.1 | 105.1 | 110.3 | 76.3 | Monolith |  |
| eGFP | N [cycles] | 400 | 90.6 | 104.8 | 109.4 | 76.2 | Monolith |  |
| S-2P | c_feed_ [USD/mg] | 6 | 32.5 | 33.0 | 74.5 | 18.2 | Monolith |  |
| S-2P | c_feed_ [USD/mg] | 15 | 45.9 | 44.9 | 90.4 | 30.5 | Beads |  |
| S-2P | c_feed_ [USD/mg] | 30 | 68.1 | 64.8 | 117.0 | 50.8 | Beads |  |
| **S-2P** | **c_feed_ [USD/mg]** | **60** | **112.7** | **104.4** | **170.2** | **91.5** | **Beads** | **yes** |
| S-2P | c_feed_ [USD/mg] | 120 | 201.7 | 183.7 | 276.6 | 172.8 | Beads |  |
| S-2P | c_lab_ [USD/h] | 15 | 104.2 | 89.9 | 139.0 | 85.8 | Beads |  |
| S-2P | c_lab_ [USD/h] | 30 | 107.0 | 94.8 | 149.4 | 87.7 | Beads |  |
| **S-2P** | **c_lab_ [USD/h]** | **60** | **112.7** | **104.4** | **170.2** | **91.5** | **Beads** | **yes** |
| S-2P | c_lab_ [USD/h] | 90 | 118.3 | 114.1 | 191.0 | 95.2 | Beads |  |
| S-2P | c_lab_ [USD/h] | 120 | 123.9 | 123.7 | 211.9 | 99.0 | Beads |  |
| S-2P | Device price | 0.5 x | 106.5 | 101.5 | 159.1 | 90.2 | Beads |  |
| S-2P | Device price | 0.75 x | 109.6 | 103.0 | 164.7 | 90.8 | Beads |  |
| **S-2P** | **Device price** | **1 x** | **112.7** | **104.4** | **170.2** | **91.5** | **Beads** | **yes** |
| S-2P | Device price | 1.25 x | 115.7 | 105.9 | 175.8 | 92.1 | Beads |  |
| S-2P | Device price | 1.5 x | 118.8 | 107.3 | 181.3 | 92.8 | Beads |  |
| S-2P | N [cycles] | 10 | 125.0 | 110.2 | 192.4 | 94.0 | Beads |  |
| **S-2P** | **N [cycles]** | **20** | **112.7** | **104.4** | **170.2** | **91.5** | **Beads** | **yes** |
| S-2P | N [cycles] | 50 | 105.2 | 100.9 | 156.9 | 89.9 | Beads |  |
| S-2P | N [cycles] | 100 | 102.8 | 99.8 | 152.5 | 89.4 | Beads |  |
| S-2P | N [cycles] | 200 | 101.5 | 99.2 | 150.3 | 89.1 | Beads |  |
| S-2P | N [cycles] | 400 | 100.9 | 98.9 | 149.2 | 89.0 | Beads |  |

**Variance-based check on the one-at-a-time design**

The sensitivity analysis reported in Table S8 varies one input at a time. That design is efficient and easy to read, but it explores only the axes through the baseline point, so on its own it cannot show whether inputs interact [[6,7]](https://www.zotero.org/google-docs/?tIGQTx). A variance-based analysis was therefore carried out as a check on whether the one-at-a-time design is adequate for Eq. (1).

In a variance-based analysis each input is treated as a random variable and the variance of the model output is apportioned between the inputs. The first-order index S(X) is the fraction of the output variance that would be removed, on average, by fixing input X at its true value. The total index ST(X) additionally includes every interaction in which X takes part, so the difference ST(X) − S(X) measures how much variance X contributes through interactions rather than on its own [[7,8]](https://www.zotero.org/google-docs/?zCeBUt). For a purely additive model the first-order indices sum to one and every difference ST(X) − S(X) is zero.

The four inputs of Table S7 were treated as independent and uniform over their ranges, and the indices were estimated with the Saltelli scheme using 2^18 = 262,144 base samples per input matrix [[6]](https://www.zotero.org/google-docs/?nAVHzo). The ranges are the same as those used for the one-at-a-time analysis, so both analyses cover the same parameter space. The uniform distributions are a device for apportioning variance across the declared ranges and are not a claim about how these costs are distributed in practice; no distribution is assigned to any input in the analysis reported in the main text.

The result is given in Table S9. Eq. (1) is additive in the mRNA feed cost, the labour and facility rate and the device price, so the only pair of inputs that can interact is the device price with the number of cycles per device, which enter the equation as a ratio. The first-order indices account for essentially all of the output variance in every matrix–construct combination, and the device price × N interaction stays below 0.2% of the variance. This confirms that the one-at-a-time design reported in Table S8 captures the behaviour of the model over these ranges, and that a sampled or variance-based design would be expected to reproduce it. The script and input data that reproduce Tables S7 to S9 and Figure S2 are deposited with the dataset (see Data availability).

**Table S9.** Variance decomposition of Eq. (1) over the ranges given in Table S7, estimated as described above. S(X) is the first-order Sobol index of input X, and the final column is the share of the output variance that the device price contributes through its interaction with N, that is ST(Pdev) − S(Pdev). All indices are dimensionless fractions of the total output variance. Because the device price and N are the only pair of inputs that can interact, the same interaction estimated from the N side gives values of comparable magnitude, at most 0.0013; both estimates sit at the level of Monte Carlo noise, which is why indices that are zero to three decimal places are occasionally returned as very small negative numbers and are reported as zero here.

| **Construct** | **Format** | **First-order index, S(c_feed_)** | **First-order index, S(c_lab_)** | **First-order index, S(P_dev_)** | **First-order index, S(N)** | **Sum of first-order indices** | **Interaction share, S_T_(P_dev_) − S(P_dev_)** |
| --- | --- | --- | --- | --- | --- | --- | --- |
| eGFP | Monolith | 0.978 | 0.009 | 0.000 | 0.004 | 0.991 | 0.0002 |
| eGFP | Beads | 0.910 | 0.108 | 0.000 | 0.001 | 1.019 | 0.0001 |
| eGFP | Membrane | 0.941 | 0.053 | 0.001 | 0.008 | 1.003 | 0.0004 |
| eGFP | Fibre | 0.986 | 0.011 | 0.000 | 0.000 | 0.998 | 0.0000 |
| S-2P | Monolith | 0.996 | 0.014 | 0.000 | 0.003 | 1.012 | 0.0008 |
| S-2P | Beads | 0.944 | 0.046 | 0.000 | 0.001 | 0.992 | 0.0000 |
| S-2P | Membrane | 0.876 | 0.115 | 0.000 | 0.008 | 0.999 | 0.0011 |
| S-2P | Fibre | 0.987 | 0.007 | 0.000 | 0.000 | 0.994 | 0.0001 |

**Table S10.** Chromatography device list prices used in the cost model. List prices are used rather than the prices actually paid, because the columns used in this study were supplied free of charge or at substantially discounted prices (see Acknowledgement), so the comparison rests on prices any purchaser could obtain.

| **Format** | **Product** | **Supplier** | **Bed volume [mL]** | **List price [USD]** | **Relative to monolith [%]** | **Catalogue no.** |
| --- | --- | --- | --- | --- | --- | --- |
| Monolith | CIMmultus Oligo-dT 18 (C12 linker) | Sartorius BIA Separations | 1.0 | 833 | 100.00 | 311.1219-2 |
| Beads | POROS GoPure Oligo (dT)25 | Thermo Fisher Scientific | 1.0 | 440 | 52.82 | A48352 |
| Membrane | Purexa OdT Maxi | Donaldson (Purilogics) | 0.2 | 500 | 60.02 | 100003046 |
| Fibre | Fibro dT adsorber | Cytiva | 0.4 | 274 | 32.89 | 17553101 |

Prices were taken from online supplier catalogues and were accurate as of 27 October 2026, converted at a EUR/USD rate of 1.11 where applicable.

**Normalised relative cost comparison**

The comparison below was carried out alongside the amortised model of Eq. (1) and gives the same ranking of the formats at the baseline. It is reported here rather than in the main text for two reasons. Its total combines categories that are each normalised to a different reference, so the total is a relative score rather than a cost share. It also holds labour and facility cost equal across formats, whereas Eq. (1) scales them with cycle time and shows that they differ by about fourfold between formats. Eq. (1) is therefore used as the cost analysis in the main text, and the figures below are given for completeness.

A relative cost analysis was performed for all four formats. The analysis combines the purchase price of the chromatography device with the efficiency with which each format converts loaded mRNA into purified product, and separates four categories: raw materials, chromatography matrix, labour, and facility and other overheads. The following assumptions were applied. (1) Every column was operated at the conditions of its near-optimal run, as given in Figure 3 and Table S5. (2) Labour and facility-associated expenditure, including cleanroom time, personnel and utilities, were taken to be constant across formats and were held at fixed shares of 4.5% and 5.5% of the reference process cost [[1,2]](https://www.zotero.org/google-docs/?R2o9oD); Eq. (1) replaces this treatment with one that scales labour and facility cost with cycle time. (3) Each format was assumed to sustain the same number of bind-elute cycles before ligand degradation or significant loss of performance. (4) Buffer compositions, system set-up and equipment usage were standardised across experiments and excluded from the variable cost comparison. (5) Device costs were taken from publicly available supplier list prices, normalised in USD, with the monolith column set to 100% (Table S10). (6) Raw-material efficiency is defined as the cost of process consumables required to obtain one milligram of purified mRNA, that is, the ratio of mRNA loaded to mRNA recovered; for each construct the format with the lowest recovery was set as the reference at 100% and the others scaled to it. The total relative cost is the sum of the four categories weighted by their shares of the reference process, 60% raw materials, 30% chromatography device, 4.5% labour and 5.5% facility and other overheads.

The results are given in Table S11. Raw-material cost tracked recovery directly. The membrane, with the lowest recovery for both constructs, set the 100% reference, while the fibre adsorber (79.1% for eGFP and 76.5% for S-2P), the beads (87.9% and 74.5%) and the monolith (94.1% and 83.7%) consumed proportionally less mRNA per milligram purified. Device cost separated the formats more sharply: on supplier prices the fibre adsorber is the least expensive device of the four at 32.9% of the monolith, followed by the beads at 52.8% and the membrane at 60.0%. Combining the two, the fibre adsorber scored lowest, at 67.3% of the monolith baseline for eGFP and 65.8% for S-2P, followed by the beads (78.6% and 70.6%), whereas the membrane (88.0% for both constructs) and the monolith (96.4% and 90.2%) scored highest. With eGFP the fibre adsorber combines the least expensive device with the highest recovery of the four formats; with S-2P its raw-material efficiency is close to that of the beads, and its advantage rests mainly on the device price.

Two limitations qualify these figures. First, assumption (3) credits every format with the same number of cycles. The monolith and the beads are sold for validated multi-cycle re-use, with at least 40 cleaning cycles demonstrated for the monolith [[3]](https://www.zotero.org/google-docs/?rkG1jH), whereas the membrane and fibre devices used here are supplied for research and process development; were the thin-bed formats operated single-use while the monolith and beads were re-used across a campaign, their device cost per batch would rise in proportion to the number of cycles. Eq. (1), which varies the number of cycles explicitly, is the appropriate treatment of this effect and is reported in Section 3.4 of the main text. Second, the raw-material category is normalised to recovery, and recovery here is bounded by the load challenge applied to each bed rather than measured at a common multiple of capacity, so that row should be read as the cost of the specific loading strategy used and not as an intrinsic property of the matrix.

**Table S11.** Normalised relative cost comparison of the four formats at the near-optimal runs, moved here from the main text. Values are percentages on the bases given in the footnotes, so the total in row (E) is a relative score rather than a cost share.

| **RNA** | **Cost category** | **Monolith column** | **Beads column** | **Membrane column** | **Fibre column** |
| --- | --- | --- | --- | --- | --- |
| eGFP | Raw-material costs [%] (A) | 94.05 | 87.88 | 100.00 | 79.11 |
| eGFP | Chromatography device costs [%] (B) | 100.00 | 52.82 | 60.02 | 32.89 |
| eGFP | Labour cost [%] (C) | 4.5 | 4.5 | 4.5 | 4.5 |
| eGFP | Facility and other costs [%] (D) | 5.5 | 5.5 | 5.5 | 5.5 |
| **eGFP** | **Total costs [%] (E)** | **96.43** | **78.57** | **88.01** | **67.33** |
| S-2P | Raw-material costs [%] (A) | 83.68 | 74.55 | 100.00 | 76.47 |
| S-2P | Chromatography device costs [%] (B) | 100.00 | 52.82 | 60.02 | 32.89 |
| S-2P | Labour cost [%] (C) | 4.5 | 4.5 | 4.5 | 4.5 |
| S-2P | Facility and other costs [%] (D) | 5.5 | 5.5 | 5.5 | 5.5 |
| **S-2P** | **Total costs [%] (E)** | **90.21** | **70.58** | **88.01** | **65.75** |

(A) The relative cost of the raw materials consumed during purification, including reagents, buffers and consumables. Values are normalised to raw-material efficiency, defined as mRNA loaded per mRNA recovered, with the least efficient format set at 100%.

(B) The relative cost of the chromatography device, taken from public supplier list prices and normalised with the monolith column at 100%. Prices and sources are given in Table S10.

(C) The cost of human resources involved in the operation of the chromatography process.

(D) Expenses related to the use of laboratory facilities, utilities and miscellaneous overheads.

(E) The weighted sum of the four categories, calculated as 0.60 x (A) + 0.30 x (B) + (C) + (D).

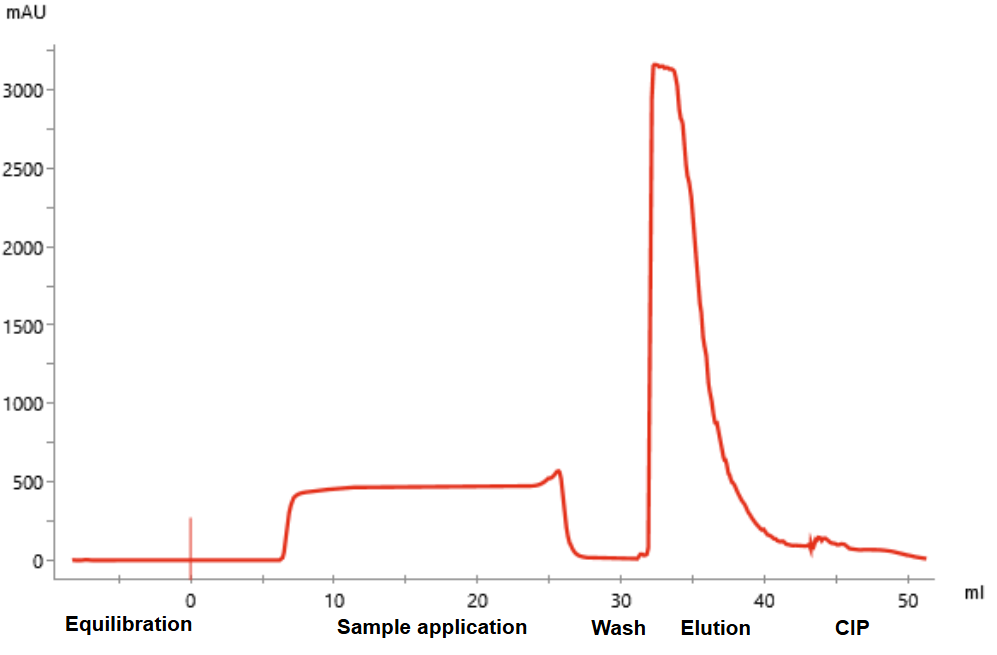

**Figure S1.** Schematic representation of a UV absorbance at 260 nm trace of a typical chromatography cycle used in the study.

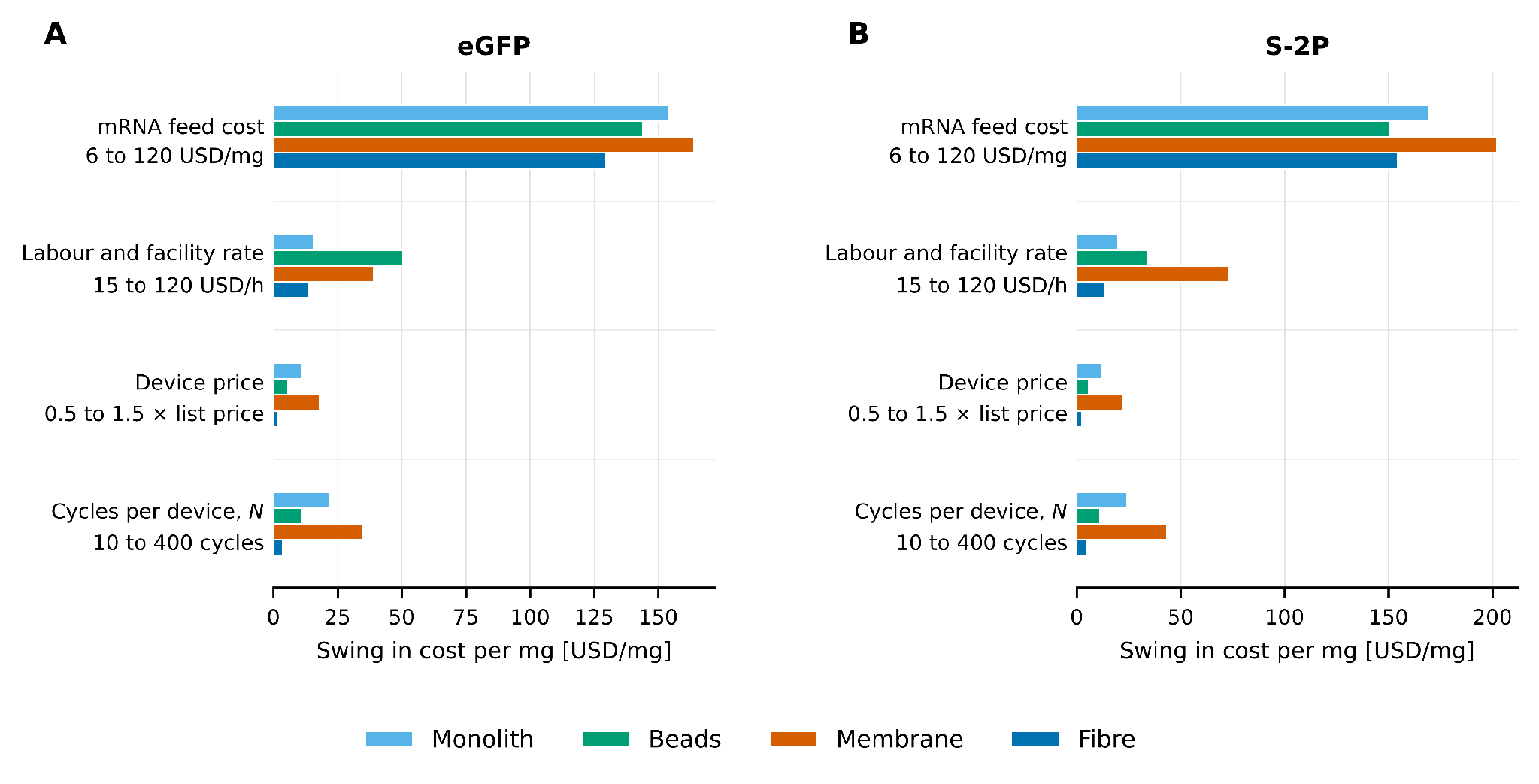

**Figure S2.** Swing in the cost per mg of purified mRNA produced by each input of Eq. (1) over the ranges in Table S7, for eGFP (A) and S-2P (B). Each bar is the difference between the cost at the two ends of that input’s range with the other three inputs held at baseline. The mRNA feed cost dominates for every format; the device price is the least influential input, and least influential of all for the fibre adsorber.

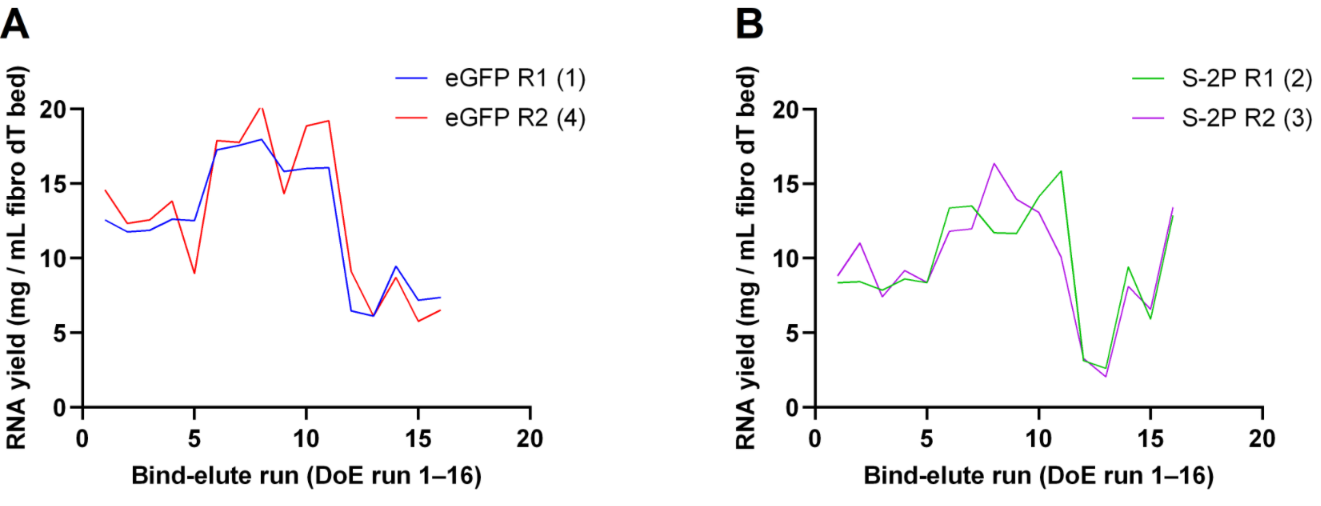

**Figure S3.** Yield stability of the 0.4 mL Fibro oligo-dT column over repeated bind–elute cycles. RNA yield (mg per mL of bed) from the 0.4 mL Fibro oligo-dT column during its lifetime study. The column completed 118 bind–elute cycles in total; the cycles plotted fall between cycles 50 and 118. Each series consists of DoE runs 1–16, which vary the Gu-HCl concentration, the load concentration and the load flow rate (factor levels in Table 2). The four series were run in sequence: eGFP R1 (1), S-2P R1 (2), S-2P R2 (3) and eGFP R2 (4). (A) eGFP mRNA, replicates R1 (blue) and R2 (red). (B) S-2P mRNA, replicates R1 (green) and R2 (purple). Within each construct, the two replicates follow the same yield pattern across the 16 conditions. eGFP R2 was the last series run, yet its yields match or exceed those of eGFP R1. This indicates no measurable loss of binding capacity over the cycles shown.
